# Widespread horizontal transfer and strong selection enhance microbial adaptation in Antarctic soils

**DOI:** 10.1101/2025.11.03.686391

**Authors:** Yongyi Peng, Laura C. Woods, Laura Perlaza-Jimenez, Rachael Lappan, Marion Jespersen, Xiyang Dong, S. Ry Holland, Steven L. Chown, Pok Man Leung, Chris Greening

**Affiliations:** Securing Antarctica’s Environmental Future, Monash University, Clayton, VIC 3800, Australia; Department of Microbiology, Biomedicine Discovery Institute, Monash University, Clayton, VIC 3800, Australia; Centre to Impact AMR, Monash University, Clayton, VIC 3800; Third Institute of Oceanography, Ministry of Natural Resources, Xiamen, Fujian, China; School of Biological Sciences, Monash University, Clayton, VIC 3800, Australia

## Abstract

Terrestrial Antarctica harbors compositionally diverse and functionally distinct microbial life. Yet the ecological and evolutionary processes enabling these communities to adapt to the polyextreme conditions of the continent remain largely unknown. Here, we address how horizontal gene transfer (HGT) and *de novo* mutations influence adaptation of microbial communities in 16 proglacial and mountainous Antarctic soils, from a combination of short- and long-read datasets. Phylogenetic reconciliation and mobile genetic element analysis of 676 metagenome-assembled genomes show that HGT events occur frequently within these microbial communities. While the transferred genes are distributed across diverse functional categories, those involved in energy metabolism are exchanged at relatively higher frequency. The genes for aerotrophy, i.e. the consumption of atmospheric trace gases to provide energy, carbon, and hydration, are among the most frequently and widely disseminated. Approximately a quarter of all carbon monoxide (CO) dehydrogenases and [NiFe]-hydrogenases that catalyze atmospheric CO and hydrogen (H_2_) oxidation are predicted to be horizontally acquired and are often closely associated with mobile genetic elements. In parallel, analysis of polymorphisms in protein-encoding genes suggests widespread purifying selection, demonstrated by a predominance of synonymous mutations. This selection is particularly intense for aerotrophy genes, providing further evidence that this process is critical for microbial survival in Antarctica. The genetic variation of hydrogenases is tightly associated with their predicted protein structures, with intense selection acting on critical sites that preserve their stability and function in Antarctic environments. Together, these findings show that previously unrecognised eco-evolutionary dynamics shape the composition and function of Antarctic desert microbial communities, and confirm aerotrophy is a strongly selected and horizontally disseminated trait.

## Introduction

Once regarded as a frozen desert with limited signs of life, Antarctica is increasingly recognized as a continent populated with metabolically active and taxonomically diverse microbial communities^1–3^. What remains unclear, however, is how this diversity arises and is maintained under the strong constraints imposed by the Antarctic environment. One potential mechanism is genetic diversification through *de novo* mutations and the acquisition of foreign DNA through horizontal gene transfer (HGT), which governs both ancient and contemporary bacterial evolution^4–6^. HGT can enable microbes to acquire novel traits that facilitate ecological fitness in extreme niches, for example introducing genes conferring advantageous phenotypes such as antibiotic resistance^7,8^, alkane metabolism^9^, metal resistance^10^, transporter functions^11^ and virulence^12^. Within populations, mutations also generate novel genotypes and serve as a fundamental driver of adaptive evolution. Yet these changes are not uniformly retained: abiotic stresses and biotic interactions impose selective pressures that favor beneficial variants while purging deleterious ones^4,13^. Thus, integrating the mechanisms of genetic variation (mutation and HGT) with natural selection offers a powerful framework for elucidating how microorganisms diversify and adapt in natural environments. Despite its relevance, we lack a systematic, community-wide understanding of HGT and selection processes in Antarctica, and the extent to which they enable adaptation to the continent’s extreme environments.

Pioneering studies in other extreme ecosystems have examined how variation processes and natural selection shape microbial adaptation. For example, surveys of microbial communities within deep-sea hydrothermal vents have revealed frequent horizontal transfer of energy metabolism and nutrient acquisition genes^14^; halophilic bacteria and archaea have been inferred to exchange genes for salt adaptation and phototrophy^15,16^; and key metabolic genes are also under strong purifying selection in deep-sea cold seeps^17–19^. By contrast, little is known about how these eco-evolutionary dynamics shape metabolic traits in Antarctica, which experiences extremes of temperature, aridity, light, UV, salinity, and nutrients. Genetic diversification is likely a particularly crucial mechanism for microbial adaptation in Antarctic soils, where microbial dispersal is restricted^20^, and survival depends on efficient use of limited energy sources. There are several reports of mobile genetic elements in Antarctic isolates^21–24^ and the possible horizontal transfer of genes associated with antimicrobial resistance, cold tolerance, and biodegradation^25–27^. Further reflecting the physicochemical pressures of the continent, there is also some evidence of cross-domain horizontal transfer of bacterial ice-binding proteins to fungi and algae in Antarctica^28,29^. Yet, the interactions among microbial communities in Antarctica remain poorly understood^30^ and HGT has not yet been characterized at a community or system-wide scale. We hypothesize that these eco-evolutionary dynamics are of particular importance in Antarctica to overcome slow growth and turnover that would otherwise result in slow rates of adaptation.

To investigate this hypothesis, we examine HGT and selection patterns of all genes encoded by microbial communities from 16 Antarctic desert sites in the Mackay Glacier region^2^. We focus primarily on aerotrophy, the major mechanism of energy acquisition in these oligotrophic environments in which atmospheric trace gases such as hydrogen (H_2_) and carbon monoxide (CO) are consumed as energy sources, often combined with fixation of carbon dioxide (CO_2_). Bacteria encode high-affinity enzymes, predominantly group 1h and 1l [NiFe]-hydrogenases and form I carbon monoxide dehydrogenases, to oxidize these gases at atmospheric levels^2,31,32^. *In situ* and *ex situ* biogeochemical measurements have demonstrated that this process can provide electrons to simultaneously support energy conservation via aerobic respiration, carbon fixation via Calvin-Benson-Bassham cycle, and hydration through metabolic water production in nutrient-poor desert ecosystems^2,31,33,34^. Based on a previous metagenomic survey of the Mackay Glacier region, bacteria that mediate this process are both numerous and diverse, comprising ~90% of the community and spanning nine phyla, including members from the dominant phyla *Actinomycetota*, *Chloroflexota*, and *Pseudomonadota*^2^. Yet it remains unclear what shapes these patterns, including whether HGT facilitates the widespread dissemination of aerotrophy genes and whether these genes are subject to selection in Antarctic environments. Here, we employ state-of-the-art methods to quantify genetic variation and selection in microbial populations and systematically analyse the gene transfer patterns, genetic variation, and selection signatures, particularly in aerotrophy genes. Through these techniques, we provide strong evidence that both combination of HGT and intense selection support microbial adaptation in Antarctic soils.

## Results and Discussion

### Widespread HGT within microbial communities is dominated by energy metabolism genes

To investigate dissemination and selective pressures within Antarctic microbial communities, we first profiled 676 medium- to high-quality metagenome-assembled genomes (MAGs) from 16 proglacial and mountainous soils. These were dereplicated from a combination of our previously generated short-read data (451 published MAGs)^2^ and newly acquired deeply sequenced long-read and short-read data (253 new MAGs) (**Tables S1 and S2**). The MAGs spanned 19 bacterial phyla and one archaeal phylum (*Thermoproteota*) and were highly represented by bacteria from *Actinomycetota* (n=215), *Pseudomonadota* (n=85), *Chloroflexota* (n=81), and *Acidobacteriota* (n=63) (**Fig. S1 and Table S2**)^2,3^. To detect HGT events, we employed MetaCHIP^35^, integrating best-match and phylogenetic reconciliation approaches, and inferred 10,942 gene transfer events involving 664 MAGs (~98.2% of the total) and 7,959 unique donor-recipient MAG pairs (**Table S3**).

Among all the phyla, *Actinomycetota* is predicted to account for over half of all inferred HGT events (53.5%), with substantial gene transfer also observed in other abundant phyla such as *Pseudomonadota*, *Chloroflexota*, *and Acidobacteriota* (**Fig. S2 and Table S4**). The obligate symbiont *Chlamydiota* exhibits the highest within-phylum MAG-pair HGT frequency (0.42; defined as the proportion of inferred HGT events relative to all possible MAG pairs), followed by *Myxococcota* (0.20), *Bacteroidota* (0.17), and *Verrucomicrobiota* (0.16) (**Fig. 1a and Table S4**). Gene exchange is predicted to occur more frequently within than between phyla, although within-phyla events may be missed due to detection difficulties. Several cross-phyla HGT events were also prominent, such as *Desulfobacterota_B* to *Acidobacteriota* (0.08) and *Deinococcota* to *Armatimonadota* (0.05). Supporting this pattern, *Desulfobacterota_B*, *Planctomycetota*, *Armatimonadota*, and *Chlamydiota* exhibited high average numbers of mobile genetic elements (MGEs) per MAG (**Fig. 1a**). HGT is often facilitated by MGEs, which may carry potentially beneficial genes and act as reservoirs of adaptive traits under environmental stress^36^. The MGE repertoire in these taxa was dominated by insertion sequences (IS) and transposons, followed by phage recombinases (**Fig. S3**), suggesting that abundant and diverse MGEs likely facilitate both intra- and inter-phyla gene flow. Overall, these findings indicate that dominant taxa and their mobilomes play a central role in shaping the HGT landscape of Antarctic microbial communities, consistent with observations of the high proportion of mobilome genes in desert soils worldwide, compared to non-dryland soils^37,38^.

**Figure 1.**
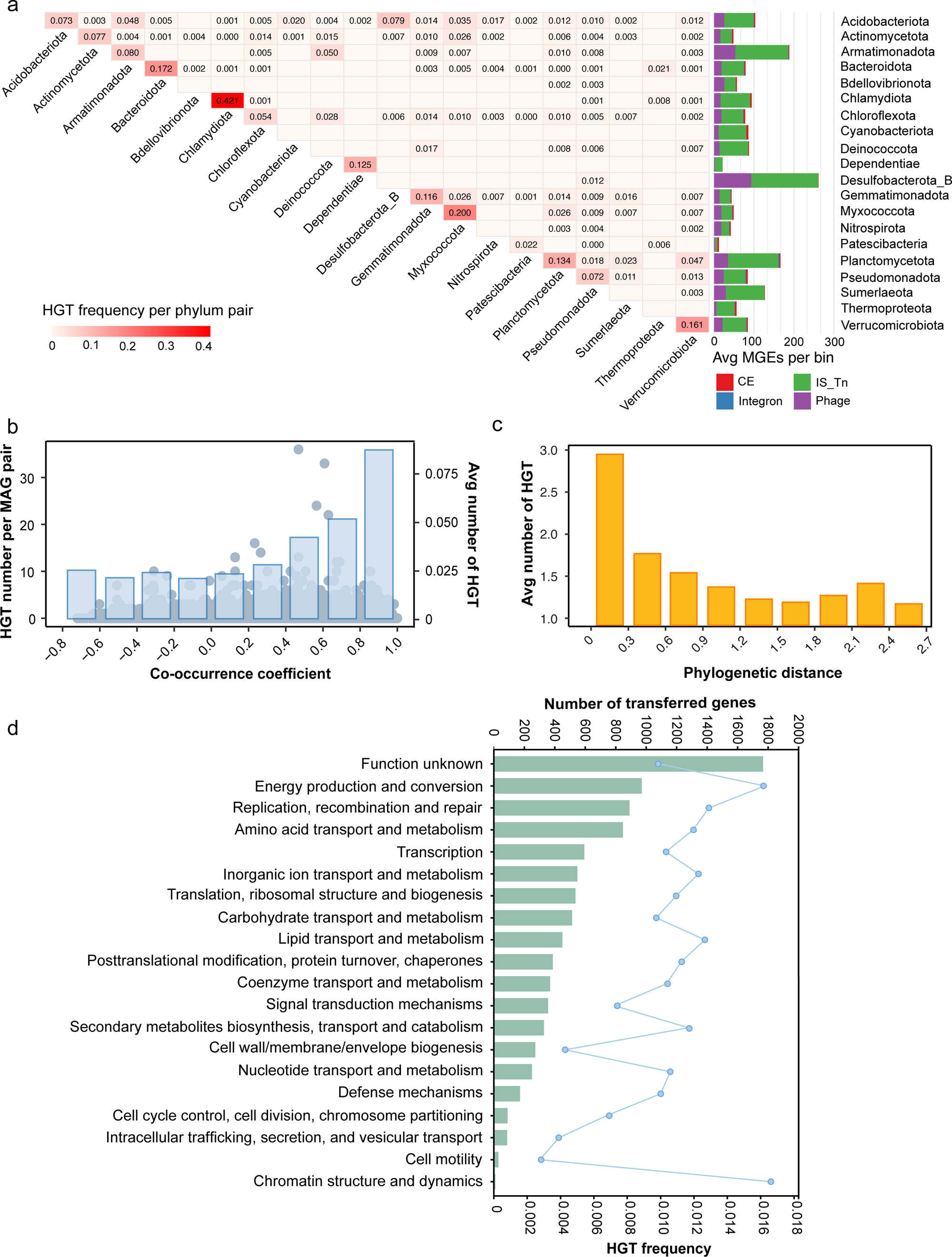
Ecological patterns of HGT across Antarctic microbial communities. (a) Community-level HGT events across different microbial phyla, indicating the HGT frequency between each phylum pair. HGT frequency is defined as the proportion of inferred HGT events relative to all possible bin pairs. The bar plot on the right shows the average number of MGEs per genome bin, with each MGE type represented by a different color. CE refers to conjugative elements and IS_Tn refers to insertion sequences (IS) and transposons. (b) Relationship between the co-occurrence coefficient and the number of HGT events. The co-occurrence coefficient represents the pairwise Spearman correlation between MAGs across samples based on their relative abundances. The scatter plot (left y-axis) displays the number of HGT events corresponding to each specific co-occurrence coefficient value for each MAG pair. The bars (right y-axis) represent the average number of HGT events within different ranges of co-occurrence coefficients. (c) Bar plot showing the average number of HGT events across different phylogenetic distances. (d) Functional distribution of horizontally transferred genes based on COG categories. The bars (top x-axis) represent the total number of transferred genes in each COG category. The line plot (bottom x-axis) shows the HGT frequency, calculated as the proportion of transferred genes relative to all genes in that category. Detailed statistics for HGT events of 676 MAGs are provided in Tables S3-7.

Both ecological interactions and evolutionary factors can influence horizontal gene transfer^39^. For Antarctic microbial communities, we studied the association between the number of HGT events per MAG and their co-occurrence patterns with other community members. As expected, we observed a positive correlation between the average number of detected transfers and co-occurrence (**Fig. 1b and Table S5**), suggesting gene transfer is more likely among microorganisms with positive co-occurrence than among those with non-significant or negative correlations. Closely related species are also more likely to transfer genes due to the generally lower phylogenetic distances, consistent with the observation that within-phylum transfers occurred more frequently than between-phylum transfers (**Fig. 1c and Table S6**). These findings are consistent with a global study of prokaryotes that revealed co-occurring species tend to exchange more genes, and that HGT frequency is also influenced by phylogenetic similarity^37,40^. We next examined the function of transferred genes to gain insights into how HGT influences microbial adaptation in Antarctic environments. Functional annotations revealed a significant enrichment of transfer events in genes associated with various metabolic functions, particularly energy production and conversion, as characterized by both a high number and high proportion of transferred genes (**Fig. 1d and Table S7**; Fisher’s exact test, P < 2.2 × 10^−16^, odds ratio = 1.70). This contrasts with a previous large-scale survey of prokaryotic genomes reporting that genes involved in metabolic processes are generally depleted in HGT events^37^, whereas energy metabolism genes in Antarctic soils have higher horizontal transfer probabilities than expected from the global dataset (Fisher’s exact test, *P* < 2.2 × 10^−16^, odds ratio = 1.40). Interdomain gene transfers from bacteria to archaea have also been shown to predominantly involve metabolic functions, suggesting such gene transfers play an important role in microbial adaptive evolution^41^. Our results indicate that strong selection in Antarctica’s polar desert soils likely favors the horizontal gene acquisition for energy conservation and resource use, potentially contributing to enhanced energetic efficiency in slow-growing, temperature-constrained, and resource-limited microbes.

### Aerotrophy genes are frequently transferred in Antarctic microbial communities

Given the enriched transfer of genes involved in energy production and conversion, we further investigated the metabolic strategies that sustain Antarctic microorganisms in extreme environments. By profiling the distribution of genes representing different energy conservation and carbon acquisition pathways, our results suggest that the prevalent taxa in oligotrophic Antarctic soils are metabolically versatile aerobes that use atmospheric trace gases to meet their energy and carbon needs. In agreement with previous findings for Antarctic polar deserts^3,31,33,42^, genes for atmospheric hydrogen and carbon monoxide oxidation, as well as carbon fixation, were widespread (**Fig. S4 and Table S8**). Genes encoding hydrogenases were present across 12 phyla and phylogenetic analyses revealed four different clades, mainly from the high-affinity group 1h and 1l uptake [NiFe]-hydrogenases. Form I CO dehydrogenases (*coxL*) were identified in 37 MAGs, all assigned to *Actinomycetota* or *Chloroflexota*. The RuBisCO large subunit gene (*rbcL*), encoding the key enzyme for carbon fixation through Calvin-Benson-Bassham cycle, was present in six dominant phyla. Of the 43 MAGs encoding Type IE RuBisCO, which exclusively mediates chemosynthetic carbon fixation, 70% also encoded high-affinity hydrogenases and 35% encoded CO dehydrogenases (**Fig. S5**). Members of Antarctic microbial communities also encoded key genes for autotrophic ammonia oxidation (*amoA*, *hbs*), sulfide oxidation (*sqr*, *fcc*) and rhodopsin-based light harvesting (*rho*) (**Fig. S4**). These results support previous findings that microbial lineages in organic carbon-deficient Antarctic soils continually harvest alternative energy sources to survive under limiting conditions.

We then surveyed the horizontal transfer of these metabolic genes between the Antarctic soil MAGs. Among the 52 profiled energy-converting metabolic genes, nine were predicted to be involved in a total of 75 horizontal transfer events across nine dominant phyla, with *Actinomycetota* accounting for the most (41 HGT events) (**Fig. 2a**). More than half of HGTs (42 events) were within-phylum transfers, further supporting the finding that closely related taxa are more likely to exchange genes. Of the metabolic genes analyzed, aerotrophy genes (including group 1h and 1l [NiFe] hydrogenases, CO dehydrogenases, and Type IE RuBisCO) each showed a transfer frequency (defined as the proportion of transferred genes relative to total gene counts) more than twice as high as that observed for other genes (**inserted in Fig. 2a**). Among them, group 1l and 1h uptake hydrogenases were the most horizontally transferred genes in these polar desert communities, with 24 HGT events involving 42 hydrogenase genes predicted across eight hydrogenase-bearing phyla (**Fig. 2a-b**). Group 1h hydrogenases are high-affinity, O_2_-insensitive enzymes capable of oxidizing atmospheric hydrogen, whereas uncharacterized group 1l hydrogenases are hypothesized to share similar O_2_ tolerance and are implicated in mixotrophic growth^43,44^. Supporting the high rates of predicted horizontal transfer, we observed significant incongruence between the hydrogenase gene tree and the corresponding species tree from the Antarctic soil MAGs (**Fig. S6**). Group 1l and 1h [NiFe] hydrogenase catalytic subunit genes were encoded by ten and nine different bacterial phyla, respectively, and the shared gene sequences displayed relatively conserved identities, ranging from 72% to 96%. HGT has previously been suggested to play a critical role in shaping the distribution of hydrogenase genes in the marine environment^45^, but has not previously been demonstrated in terrestrial context. The transfer of highly similar sequences may indicate that high-affinity hydrogenase functions are highly conserved and important for adaptation to the Antarctic environment.

**Figure 2.**
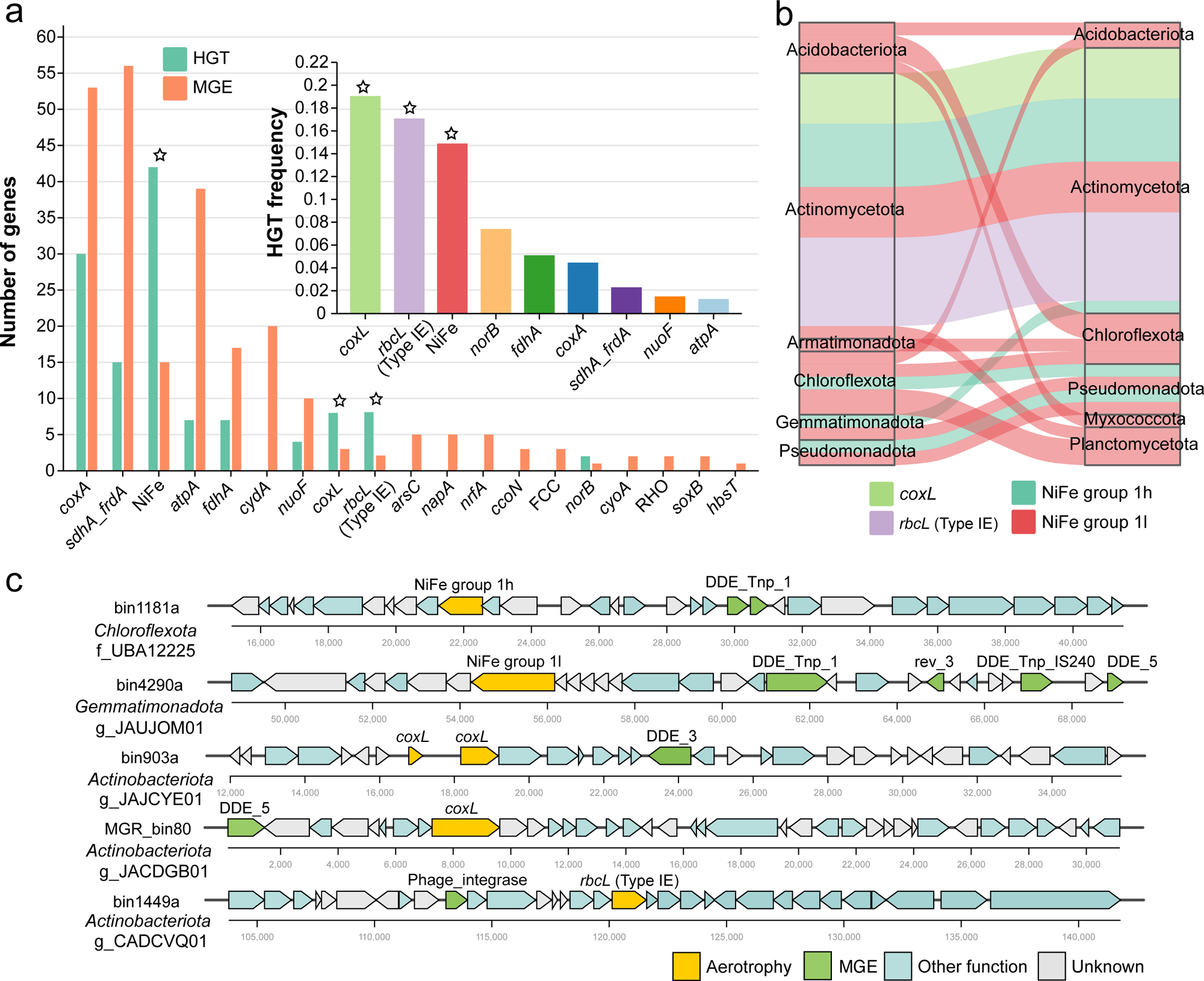
Aerotrophy genes are transferred across diverse bacterial phyla and subject to mobile genetic element transfers. (a) Comparison of gene numbers associated with HGT (green) and MGEs (orange) for key metabolic marker genes involved in energy conservation. The inset displays the HGT frequency of the top transferred genes, calculated as the proportion of transferred genes among all identified genes for each type. Aerotrophy genes are indicated by asterisks above the bars. (b) Predicted gene flow of aerotrophy genes within Antarctic microbial communities. Bands connect donors and recipients, with the width of the band correlating to the number of HGTs and the colour corresponding to the gene types. (c) Representative gene clusters containing aerotrophy genes and MGEs. Genes are color-coded by functional category: yellow, aerotrophy genes; green, MGEs; blue, other functional genes; grey, unknown function. Detailed statistics for HGT events of aerotrophy genes and MGEs are provided in Tables S8 and 9.

Among the metabolic genes analysed for HGT events between the Antarctic MAGs, genes mediating succinate oxidation (*sdhA*), aerobic respiration (*coxA*), and ATP synthase (*atpA*) were most frequently co-localized with MGEs at the genomic level (i.e., within 10 kb up- or down-stream) (**Fig. 2a, Fig. S7 and Table S9**). Hydrogenases were also frequently genomically associated with MGEs, especially DDE-type recombinases (12 of 18 MGEs), which may facilitate their inter- or intra-phylum transfer (**Figure 2c**). These findings further support that MGE-mediated transfer facilitates the dissemination of key metabolic functions in Antarctic soils, thereby enhancing community-wide capacity to cope with energy limitations and environmental stress.

### Aerotrophy genes are highly genetically conserved and under strong purifying selection

To explore the selection pressure acting on these metabolic genes, we assessed the genetic variability and their selection based on metagenomic reads. A total of 13,277,425 single-codon variants (SCVs) of all genes were identified within 676 genomes, corresponding to 829,839 SCVs per metagenome (n = 16) on average. SCVs were further classified as synonymous (s) or nonsynonymous mutations (ns), and the synonymous and nonsynonymous substitution rates were calculated at each position within genes, with the ratio expressed as the pN/pS value. pN/pS^(gene)^ is a metric that assesses whether genes are under purifying selection (pN/pS^(gene)^ < 1, the selective removal of deleterious mutations) or positive selection (pN/pS^(gene)^ > 1, the fixation of advantageous mutations)^46,47^. Overall, widespread purifying selection was observed in Antarctic protein-encoding genes (median pN/pS^(gene)^ = 0.17, ratio of pS^(site)^ to pN^(site)^ = 6:1).

Aerotrophy genes exhibited significantly lower pN/pS^(gene)^ values (0.09 for [NiFe]-hydrogenase, 0.12 for CO dehydrogenase and RuBisCO catalytic subunit genes on average) compared to the sample-average pN/pS value of genomes (0.22), confirming the elevated strength of natural selection acting on these genes in Antarctic soils (**Fig. 3**). In particular, SCVs in high-affinity hydrogenase genes were overwhelmingly synonymous, with pS^(site)^ exceeding pN^(site)^ by a ratio of 12:1 (**Fig. 3 and Table S10**). Furthermore, group 1l hydrogenases exhibited significantly lower pN/pS^(gene)^ values than group 1h, suggesting distinct evolutionary trajectories for each hydrogenase group, with the more prevalent group 1l genes subject to intensified purifying selection. Such strong purifying selection of Antarctic hydrogenases is consistent with expectations, given the importance of H_2_ oxidation for microbial survival and energy acquisition in Antarctic desert soils.

**Figure 3.**
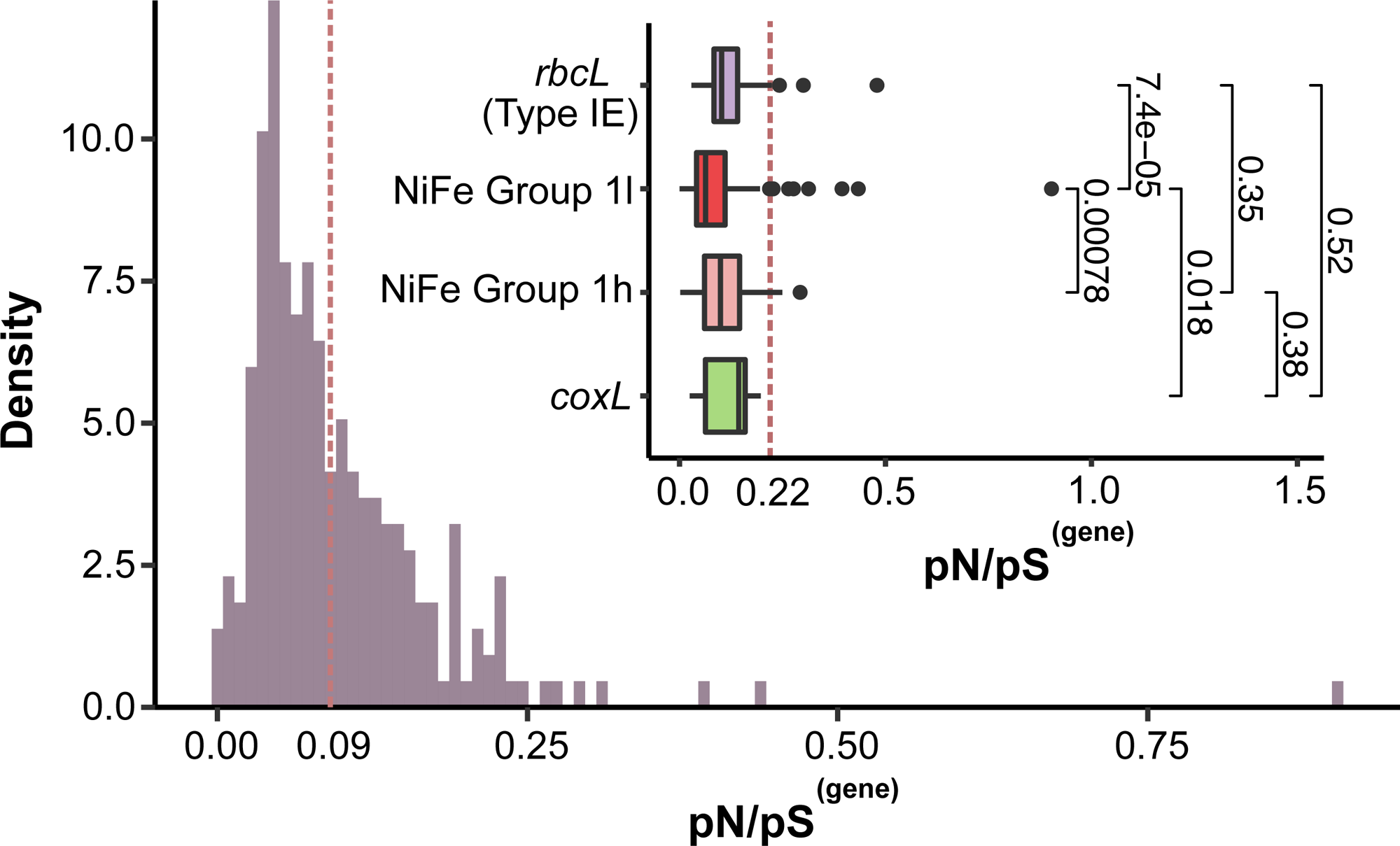
Selection pressures of aerotrophy genes in Antarctic soils. Distributions of pN/pS^(gene)^ for hydrogenase genes across samples. The dashed line indicates the sample-average pN/pS^(gene)^ for hydrogenase genes (0.09). The inset compares pN/pS^(gene)^ across two groups of hydrogenases, CO dehydrogenases, and Type IE RuBisCO. P-value of differences as calculated using a Wilcoxon rank-sum test. The dashed line in the insert indicates the overall average pN/pS^(gene)^ of genomes containing hydrogenases (0.22), highlighting the relatively low ratio in these specific gene families. Source data is available in Table S10.

Significant differences of pN/pS^(gene)^ among microbial phyla were also observed (*P* < 0.001), indicating that different Antarctic taxa likely experience varying evolutionary constraints (**Fig. S8a**). Furthermore, although significantly more pN/pS^(gene)^ variation occurred between genes compared to between samples (explaining 73.6% versus 1.6% of pN/pS^(gene)^ variation, ANOVA), the pN/pS^(gene)^ still varied significantly from sample to sample (*P* < 0.001; **Fig. S8b**). This suggests that selection strength varies in response to the range of environmental properties observed across samples. Thus, although aerotrophy genes are universally under strong selection in Antarctica, subtle differences in selection strength exist across samples and phyla, highlighting the environmental dependency of evolutionary processes in microbial communities, as also observed in other microbial systems^17,18,46^.

### Intense purification of structurally and functionally critical sites within hydrogenases

We next investigated site-specific selection within group 1h and 1l hydrogenases by integrating polymorphism rates with protein biophysical characteristics, following a published approach^46^. Within the predicted structures of hydrogenase catalytic subunits, relative solvent accessibility (RSA) and distance to ligand (DTL) for each residue were calculated to quantify structural and functional constraints (see **Methods**). RSA (ranging from 0 to 1) reflects the extent of site exposure, while DTL measures the distance to the nearest predicted ligand-binding site^46^. By comparing the distributions of synonymous polymorphism (pS^(site)^) and nonsynonymous polymorphism (pN^(site)^) rates relative to biophysical properties of hydrogenase structures, we found that pN^(site)^ exhibited a strong preference for sites with higher RSA and DTL values than pS^(site)^ (**Fig. 4a-b**). The average per-site pN^(site)^ throughout the hydrogenases was 9.0 × 10^−4^, markedly dropping to 6.8 × 10^−4^ at predicted buried sites (RSA = 0) and 6.8 × 10^−5^ at ligand-binding sites (DTL = 0). These patterns suggest that buried sites and ligand-binding sites are subject to purifying nonsynonymous changes while remaining relatively tolerant to synonymous changes. This is consistent with our structural studies of high-affinity hydrogenases (i.e. the group 2a [NiFe]-hydrogenase Huc), showing they are tightly packed enzymes with inflexible metal clusters and narrow gas channels^32^, in which amino acid substitutions in most position would likely to disrupt enzyme function. Such intolerance to nonsynonymous changes at key functional sites further supports the critical importance of these enzymes to Antarctic desert soil microbiota.

**Figure 4.**
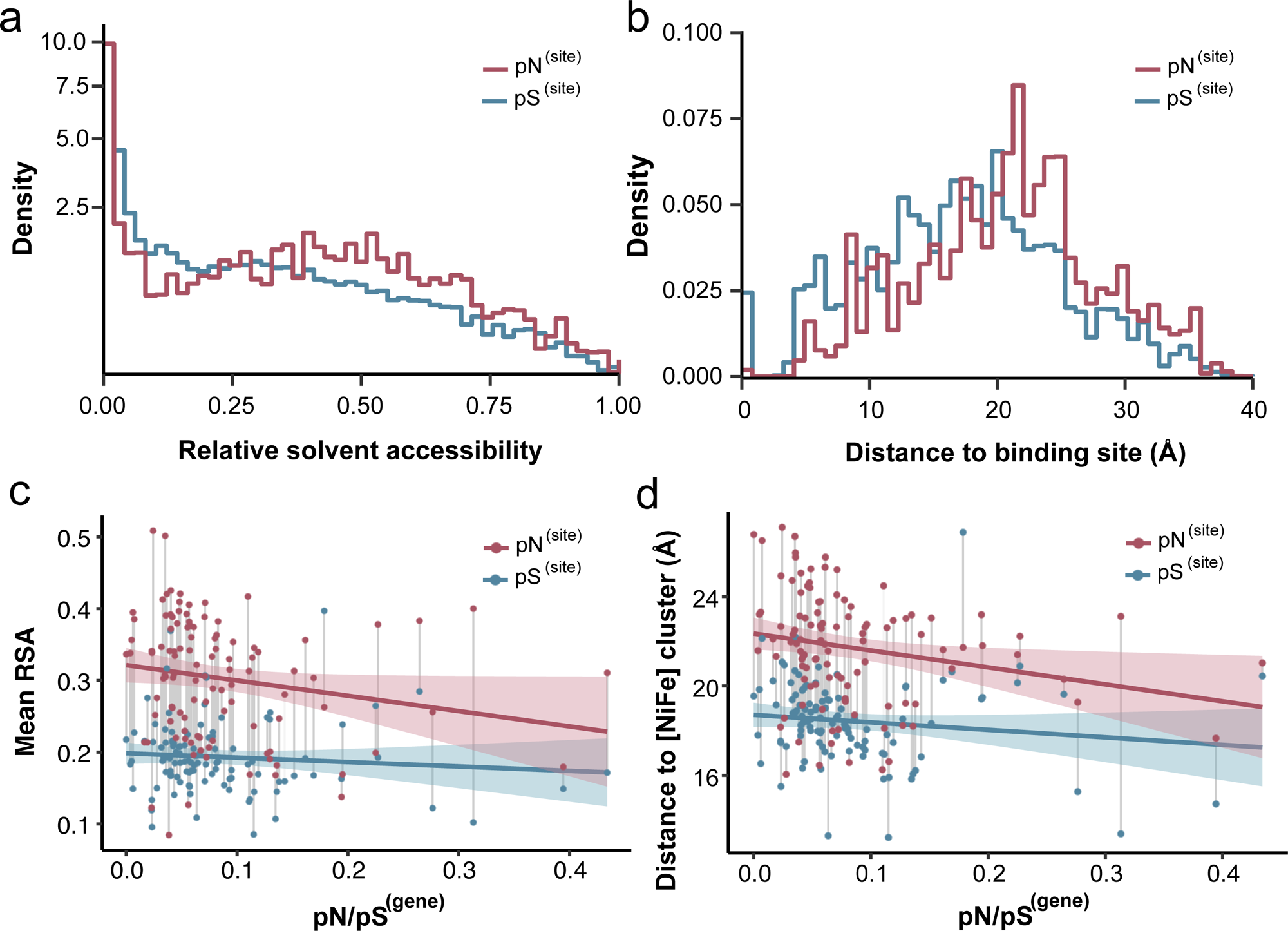
Functional and structural constraints limit nonsynonymous variations in hydrogenase genes. (a-b) Distributions of pN^(site)^ (red) and pS^(site)^ (blue) for [NiFe] hydrogenase genes in relation to relative solvent accessibility (RSA) and distance to ligand (DTL). RSA and DTL were calculated from monomeric hydrogenase structures. (c-d) Correlation between pN/pS^(gene)^ and average RSA (c) or DTL (d) across gene-sample pair. Average RSA and DTL values were calculated by weighting site-level values using pN^(site)^ and pS^(site)^. RSA and DTL values were derived from the hydrogenase complex structure, with DTL in (d) representing the distance to [NiFe]-catalytic center. Detailed statistics for pN^(site)^ and pS^(site)^ of hydrogenase genes are available in Table S11.

[NiFe]-hydrogenases function as multisubunit complexes^32^, so the RSA and DTL of individual sites may differ once assembled into the quaternary structure. To address this, we focused on group 1l hydrogenases, the most prevalent group in the sampled soils, to further investigate the relationship between polymorphism and selective processes within the complexes. Predicted quaternary structure of the group 1l [NiFe]-hydrogenase from the experimentally validated atmospheric H_2_ oxidizer *Hymenobacter roseosalivarius*^2^ shows the typical subunit composition. The large subunit (HylL) harbors the [NiFe] active site and the small subunit (HylS) contains three [4Fe-4S] clusters, forming a stable (HylLS)_2_ dimer (**Fig. 5a, right**), similar to the group 1h hydrogenases^48^. In addition, AlphaFold modelling predicted five transmembrane α-helices (HylTM1-5) that intertwine to form two tube-like structures that scaffold the cytosolic catalytic core to the membrane (**Fig. 5a, bottom left**), potentially allowing direct electron transfer into the quinone pool (**Fig. S9**). This arrangement represents a distinct orientation compared with earlier predictions based solely on sequence analysis^2^. The transmembrane helices adjacent to the C-terminal extension of HylS are highly hydrophobic and electrostatically neutral, consistent with their being membrane-embedded regions (**Fig. S10**). After superposing the predicted large subunits of Hyl from *H. roseosalivarius* and from a member of the *Sporichthyaceae* family (rmsd of 0.586 Å for 529-Cα aligned; **Fig. 5a, top left**), we recalculated RSA and DTL values, focusing on the distance to the bimetallic catalytic cluster of the enzyme^48^, which is structurally conserved in all [NiFe]-hydrogenases, including the most structurally similar resolved enzyme, the group 1h [NiFe]-hydrogenase (PDB: 5AA5, rmsd of 0.83 Å for 499-Cα aligned) (**Fig. S11**)^48^.

**Figure 5.**
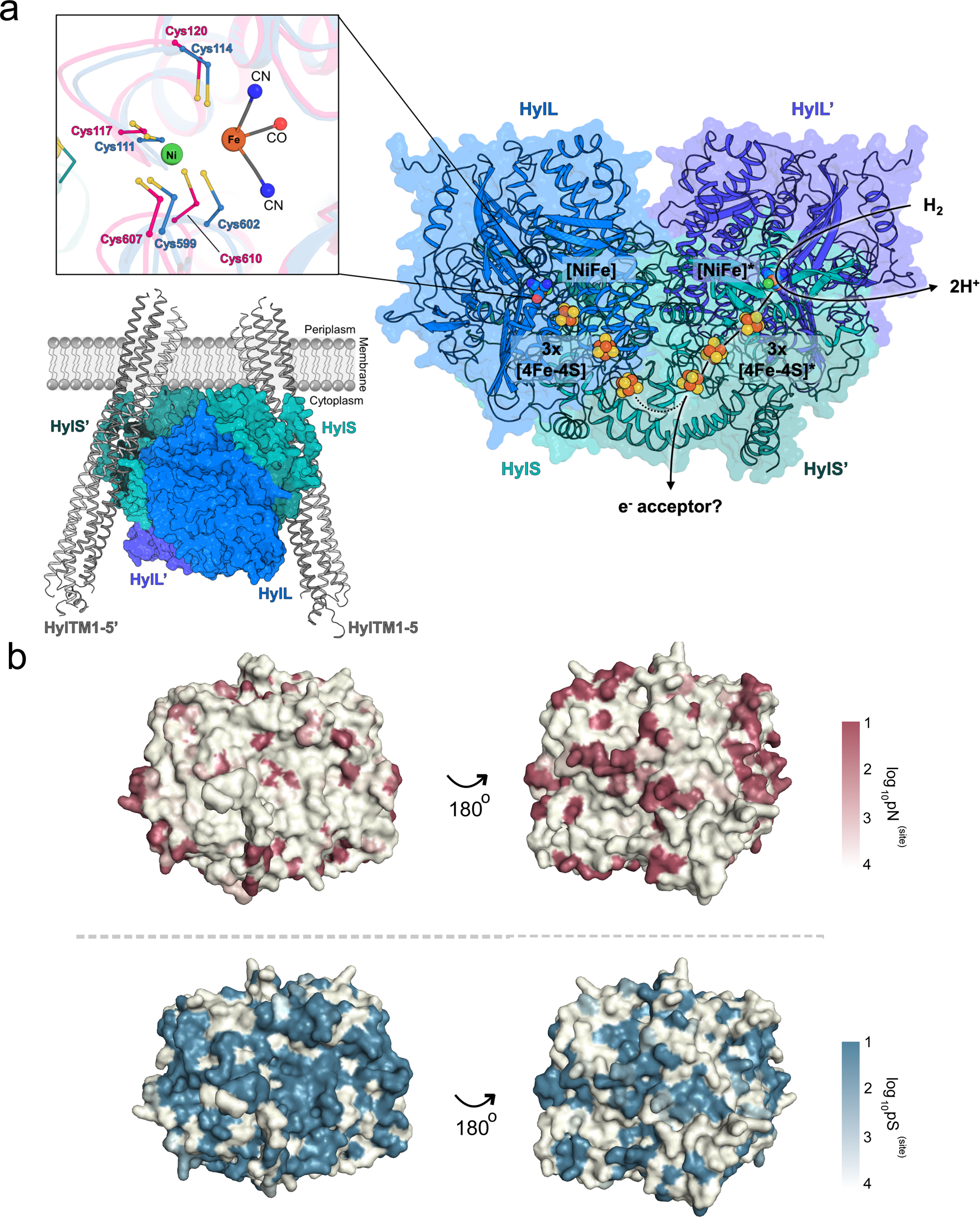
Polymorphism patterns in group 1l [NiFe] hydrogenases. (a) Right: AlphaFold3-predicted structure of the Group 1l [NiFe]-hydrogenase from *Hymenobacter roseosalivarius*. The enzyme is shown in surface and cartoon representation, with the large subunit (HylL) colored in blue and the small subunit (HylS) in teal. A star (*) marks the subunits and metal clusters of the second heterodimer, where the large subunit is shown in purple-blue and the small subunit in dark teal. The [NiFe] active site and the [Fe-S] clusters are shown as spheres. Top left: Superposition of the [NiFe] active sites from the predicted models of the *H. roseosalivarius* Group 1l [NiFe]-hydrogenase (blue) and that of an *Acidothermales* bacterium (*Sporichthyaceae* family, pink). Nitrogen, oxygen, sulfur, iron and nickel atoms are colored in blue, red, yellow, brown and green, respectively. Bottom left: Predicted membrane association of the *H. roseosalivarius* Group 1l [NiFe]-hydrogenase. The dimer is shown in surface representation, while the transmembrane helices (HylTM1-5 and HylTM1′-5′) are depicted in grey cartoon forming tube-like scaffolds that anchor the cytosolic catalytic core to the membrane. This arrangement positions the distal [4Fe-4S] clusters of the small subunits near the transmembrane domains, potentially providing a structural basis for electron transfer into the quinone pool. (b) Predicted monomeric structures of the group 1l [NiFe]-hydrogenase large subunit from *Sporichthyaceae*. Surface color indicates sample-averaged log_10_-transformed pN^(site)^ (red, upper panel) and pS^(site)^ (blue, lower panel) values at each residue, with darker colors indicating higher substitution rates. Left panels show residues that are more buried and functionally constrained, while right panels highlight more exposed and less constrained regions.

The clear shift in pN^(site)^ toward high RSA and DTL sites was observed for hydrogenase catalytic subunits in the complex, such as the highly conserved active site-coordinating cysteine residues (**Fig. 4c-d and Fig. S12**). Averaged across samples, the mean RSA value was higher for ns-polymorphisms (0.30) compared to s-polymorphisms (0.19), and a similar pattern was observed for the mean DTL (21.7 Å for ns-polymorphisms and 18.4 Å for s-polymorphisms). These findings align well with the previous observations^46,49^, for example in yeast metabolic proteins, where small-molecule-binding sites are evolving under selective constraints compared to surface residues. For ns-polymorphisms, the distance from sites to the bimetallic catalytic cluster varied markedly and was profoundly influenced by sample-average pN/pS^(gene)^. The average pN^(site)^ at active site-coordinating cysteine residues in the large subunits was markedly lower than the overall per-site average across the group 1l hydrogenases (9.5 × 10^−5^ versus 7.3 × 10^−4^), highlighting that residues predicted to play critical structural roles are indeed highly conserved. To further investigate the tight association between the per-site polymorphism rates and estimated protein structure properties, we visualized the distribution of sample-average pN^(site)^ and pS^(site)^ across the protein (**Fig. 5b**). While s-polymorphisms were more evenly distributed across the predicted subunit structure, ns-polymorphisms were concentrated in structurally flexible or surface-exposed regions where changes might be less detrimental to protein functioning. Consistent with these findings, highly conserved amino acid positions in the hydrogenase protein tend to constrain nonsynonymous variation (R^2^ = 0.074, *P* < 0.001; **Fig. S12 and Table S12**). These observations highlight that the extreme Antarctic environment exerts strong selection for key metabolic enzymes, with increased elimination of nonsynonymous mutations at buried and functional constraint sites.

## Conclusion

This work provides evidence that two key evolutionary processes shape microbial adaptation to environmental extremes in terrestrial Antarctica: HGT and strong purifying selection. HGT events are widespread within Antarctic microbial communities, as demonstrated through in-depth analysis of metabolically important genes for aerotrophy, conferring selective advantages for energetic efficiency and resource acquisition. This not only highlights aerotrophy as a fundamental process sustaining communities in nutrient-depleted polar desert soils, as others have shown^2,31,33,50^, but provides new insights into how Antarctic microbial communities actively evolve through horizontal inheritance, enabling genetic diversification even when cell duplication is limiting. Our results suggest that aerotrophy genes are closer to “hot” gene flow islands, evolutionarily independent from the core genome, due to their adaptive ecological roles within niches^51^. This echoes patterns of genetic transmission observed in other extreme^52^ or marine^53,54^ environments, but not previously reported in Antarctica. In dispersal-limited, slow-growing systems, such gene-sharing networks represent a crucial source of evolutionary innovation. In parallel, we also observed strong signals of purifying selection that act to preserve functions of core catalytic gene functions of widely dispersed genes, while permitting subtle changes that enhance diversity, as revealed by the predominance of synonymous mutations. This pressure is significantly intensified for genes mediating aerotrophy, with protein structural analysis showing an even higher conservation of critical residues near active sites and buried sites of the ubiquitous hydrogenase enzymes. Together, these forces result in the broad distribution and functional conservation of aerotrophy genes, such as high-affinity [NiFe]-hydrogenases, across diverse microbial lineages.

Our identification of key eco-evolutionary strategies underpinning microbial diversity and adaptation in Antarctica address a key eco-evolutionary shortfall identified for the region^30^. This study provides critical evidence that Antarctic microbial communities are not static or inactive, as once thought, but rather continually metabolising and genetically adapting to persist and grow under polyextreme conditions. With this understanding, Antarctica communities may be already poised to respond to a changing climate, as widespread HGT may enable rapid diversification in the face of environmental change. Altogether, these findings underscore that hidden eco-evolutionary dynamics are central for maintaining both the diversity and productivity of Antarctic soil microbes in the current climate, but whether they increase resilience or vulnerability under future changes remains to be understood.

## Methods

### Short- and long-read metagenomic sequencing

Metagenomes were sequenced from 16 proglacial and mountainous soil samples collected from north of the Mackay Glacier region in South Victoria Land, Antarctica (January 2015). Details of sample collection, DNA sequencing, and metagenomic analysis for the 16 soil samples are available in our previous work^2,7,55^. A total of 451 MAGs were derived from the previous 16 Illumina metagenomes. For this study, additional short- and long-read sequencing was performed for one soil sample from Mount Seuss (MS6-5).

MS6-5 soil sample was obtained and placed in sterile 50 ml Falcon tubes, kept below 0 °C during transport, and stored at −80 °C in the laboratory, as previously described^2^. DNA was extracted from a total of 9 g of soil (0.5 g per extract) using the FastDNA Spin Kit for Soil (MP Biomedicals) as per manufacturer’s instructions. Homogenisation was performed on a Precellys 24 instrument (Bertin Technologies) at 5,000 rpm for 40 seconds, and DNA eluted in 50 μl nuclease-free water followed by re-elution with the first eluate to maximise yield. All extracts were then pooled to provide sufficient DNA and volume for all downstream approaches.

The DNA extracts from MS6-5 were sequenced through three technologies. For short-read sequencing, BGI paired-end sequencing (2 × 100 bp) was performed using the DNBseq PE100 platform (BGI-Hong Kong), producing 27 Gb of raw reads. In addition, Illumina paired-end sequencing (2 × 150 bp) on the NextSeq500 platform at the Australian Centre for Ecogenomics (ACE) yielded a further 62 Gb of raw data. For the long-read data, genomic DNA was submitted to KeyGene (Wageningen, the Netherlands) for Oxford Nanopore sequencing. Quality control included DNA quantification and size assessment using the Femto Pulse system. A total of 869 ng of purified DNA was used for library preparation with the SQK-LSK110 kit and sequenced on a FLO-PRO002 (R9.4.1) flow cell using the PromethION P24 platform over 72 hours. Basecalling with the high-accuracy model (v20.06.18) yielded ~25.6 million reads (Q > 7), totalling ~141.8 Gb of data.

### Metagenome hybrid assembly and binning

Short-read data metagenomic analysis followed our established workflow, including quality control with BBTools v38.80 and assembly using metaSPAdes v3.13.1 ^56^. Long-read sequencing data were assembled using metaFlye v2.8^57^ and subsequently polished with Racon^58^. A hybrid assembly combining both long-read and short-read data was performed using the hybrid mode in metaSPAdes^56^.

Genome binning was performed using multiple binning tools, including MetaBAT2 v2.15^59^, MaxBin2 v2.1.1^60^, CONCOCT v1.1.0^61^, and VAMB v3.0.2^62^, and bins from the same assembly were dereplicated using the DAS_Tool v1.1.6^63^. To maximise MAG recovery, we additionally performed iterative subtractive binning as described by Rodriguez et al. (2020), which leverages read subsampling to help the assembler perform better assembly graphs and polish the final bins. In brief, reads that did not map to the initial bins were extracted and used for repeated rounds of assembly and binning using the same procedures described above. Finally, all bins were pooled across different assemblies and dereplicated and quality-filtered using dRep v3.4.3 (with an ANI ≥ 99%)^64^ and CheckM v1.1.2 (completeness > 50%, contamination < 10%)^65^, yielding a total of 253 bins. A final set of 676 MAGs was obtained by dereplicating these 253 newly generated bins along with 451 previously identified MAGs, all with an ANI of 99%. Taxonomic classifications of the resulting 676 MAGs were assigned using GTDB-Tk v2.4.0^66^ with the Genome Taxonomy Database (release 220)^67^. The phylogenomic tree for MAGs was inferred based on concatenation of 43 conserved single-copy genes that were used as phylogenetic markers in CheckM v1.1.2^65^, constructed using IQ-TREE v2.3.6 with best-fit model and 1000 ultrafast bootstraps^68^. Distances between genome pairs were extracted from the phylogenomic trees using the dist function in ETE Toolkit v.3^69^. Open reading frames from each MAG were predicted by Prodigal v.2.6.3^70^.

### Functional annotations

For metabolic annotation, we screened the predicted genes of derived MAGs against custom protein databases of representative metabolic marker genes^71^ using DIAMOND v2.1.10 (query cover > 80%)^72^. The metabolic markers included genes involved in sulfur cycling, nitrogen cycling, iron cycling, reductive dehalogenation, phototrophy, methane cycling, hydrogen cycling (large subunit of [NiFe]-, [FeFe]-, and [Fe]-hydrogenases), isoprene oxidation, carbon monoxide oxidation, succinate oxidation, fumarate reduction, carbon fixation, oxidative phosphorylation, NADH oxidation, aerobic respiration, formate oxidation, arsenic cycling, and selenium cycling. Results were filtered based on an identity threshold of 50%, except for group 4 [NiFe]-hydrogenases, [FeFe]-hydrogenases, CoxL, AmoA, NuoF, RbcL, MmoA, PsbA, AtpA and NxrA (all 60%), PsaA (80%), PsbA, IsoA, ARO and YgfK (70%), HbsT (65%), RdhA (45%), Cyc2 (35%) and energy-converting rhodopsin (30%).

For these genes, sequences with length less than 20% of the average were subjected to further inspection using two approaches: structural prediction and phylogenetic analysis. Three-dimensional structures were predicted using AlphaFold v2.0 with the full_dbs database setting^73^. Reference protein structures were then collected from the RCSB PDB database when experimentally resolved structures were available; otherwise, AlphaFold-predicted models of reviewed reference proteins were retrieved from UniProt. Structural comparisons with reference proteins were performed using Foldseek (v8.ef4e960), employing both 3Di and amino acid-based alignments^74^. Only protein structures passing the threshold defined by the Foldseek easy-search module (TM-score > 0.3, e-value < 0.001) were retained for downstream analyses. In parallel, phylogenetic analyses were performed to validate the clades of aerotrophy genes ([NiFe]-hydrogenases, CoxL, RbcL) and their reference sequences based on amino acid alignments. Sequences were aligned using MUSCLE v3.8.1551^75^ and trimmed with TrimAL v1.5.0^76^ using default parameters. Maximum-likelihood trees were generated with IQ-TREE v2.3.6 using the -m MFP -B 1000 options^68^, and all trees were visualized using iTOL^77^.

### Structural analysis of hydrogenases

Protein complex structure of group 1l [NiFe]-hydrogenase was predicted using AlphaFold3 on AlphaFold Server, enabling accurate modeling of intermolecular interactions^78^. Structures were visualized and exported as images using PyMOL v3.1.4 (http://www.pymol.org)^79^.

Hydrophobicity and electrostatic surface analysis of structural complex models were performed using UCSF Chimera v1.19^80^. Hydrophobicity values for amino acid residues were assigned automatically based on the Kyte-Doolittle scale. Electrostatic potential surfaces were calculated using Coulomb’s law as implemented in Chimera.

The conserved and variable regions of hydrogenase proteins were analyzed using ConSurf server^81^. Briefly, homologous sequences of the amino acid sequence were identified using HMMER against the UniRef90 database (E-value cutoff: 0.0001)^82^. These sequences were deduplicated using CD-HIT ^83^ and then aligned using MAFFT. A phylogenetic tree was constructed using the neighbor-joining algorithm implemented in the Rate4Site program^84^. Position-specific conservation scores were calculated using the Bayesian method and subsequently mapped onto the protein sequence and visualized on the 3D structure.

### HGT analysis

For HGT event detection, MetaCHIP v1.10.13 was employed to identify horizontal gene transfer within microbial MAGs through the combination of best-match and phylogenetic approaches^35^. Input genomes were first grouped according to their taxonomic classifications at multiple ranks (phylum, class, order, family and genus). In the best-match approach, open reading frames (ORFs) of all MAGs were first predicted using Prodigal, followed by an all-against-all BLASTN search of the predicted ORFs, applying a minimum alignment length of 200 bp, at least 75% coverage, a 90% sequence identity, and an 80% end-match identity. Genes showing higher identity to non-self groups than to self-group were considered HGT candidates, and only those in the top 10% of identity were retained. To validate and infer the direction of putative HGTs, a phylogenetic reconciliation approach was applied. Protein trees for each candidate HGT pair were constructed using MAFFT^85^ and FastTree v2.1.11^86^. A corresponding species tree was generated using 43 universal single-copy genes (SCGs) as defined by CheckM^65^. Gene and species trees were then reconciled using Ranger-DTL v2.0^87^ to identify duplication, transfer, and loss events.

To conduct functional enrichment and MGE analysis, further genome annotation was performed. The amino acid sequences from 676 MAGs were functionally annotated using eggNOG-mapper (v2.1.9; default parameters) for eggNOG 5.0^88,89^, and each gene was classified into the corresponding Clusters of Orthologous Genes (COG) categories. Genes associated with the mobilome, including integrons, integrative conjugative elements (ICEs), insertion sequence (IS) elements, and transposons, were identified by performing HMM searches against the 68 marker HMM profiles provided by proMGE (http://promge.embl.de/)^90^.

### Abundance and co-occurrence analysis

To explore factors influencing horizontal gene transfer, we calculated the relative abundance and co-occurrence patterns of MAGs. Relative abundance was determined using CoverM^91^ in genome mode (v0.6.0), with parameters set to -min-read-percent-identity 0.95, -min-read-aligned-percent 0.75, -trim-min 0.10, and -trim-max 0.90. MAGs with relative abundances greater than 0.01% and detected in more than one of the five analysed samples were included in downstream analyses. Microbial co-occurrence network analysis was performed using the R packages igraph^92^ and WGCNA^93^. Spearman correlation coefficients were computed for pairwise MAG associations, and statistical significance was assessed using False Discovery Rate (FDR) correction of P-values.

### Calculation of evolutionary metrics

The polymorphisms of protein-encoding genes were examined using Anvi’o’s framework for microbial population genetics. Filtered reads from 16 soil samples were mapped to all MAGs using Bowtie2. A contigs database of these MAGs was then generated in Anvi’o (v8)^46^ using ‘anvi-gen-contigs-database’, incorporating ORFs previously predicted by MetaCHIP v1.10.13 via the ‘--external-gene-calls’ parameter. Nucleotide-level metrics, including coverage, single nucleotide variants (SNVs), single amino acid variants (SAAVs) and single codon variants (SCVs), were calculated from the read mappings using Anvi’o profile module. Only SNV positions with > 10× coverage were retained. The synonymous (s) and nonsynonymous (ns) polymorphism rates per residue (pS^(site)^ and pN^(site)^) were calculated and filtered to include only variants with at least 0.04 departure from the consensus and a minimum coverage of 20×.

Hydrogenase genes with high-quality predicted structure (pLDDT > 80) were selected for structure-based polymorphism analyses. A structure database for these high-quality hydrogenase protein structures was created using ‘anvi-gen-structure-database’. For each residue, the relative solvent accessibility (RSA) was computed within Anvi’o using the DSSP module^94^, which quantifies the ratio of the observed solvent-accessible surface area (ASA) of a residue to its theoretical maximum accessibility in an extended tripeptide conformation. RSA values thus range from 0 (completely buried) to 1 (fully exposed), reflecting the degree of residue exposure to solvent. Ligand-binding residues were predicted using InteracDome^95^ by scanning domain-ligand interaction profiles across Pfam-annotated domains, and residues with per-residue binding frequencies > 0.5 were considered ligand-interacting. The distance to ligand (DTL) was calculated using append_dist_to_lig.py provided by Anvi’o (https://merenlab.org/data/anvio-structure/chapter-III/). Briefly, this script loads 3D atomic coordinates from the structure database and computes the minimum center-of-mass (COM) Euclidean distance between each residue and all predicted ligand-binding residues within the same protein. The key active site-coordinating cysteine residues of hydrogenases were further identified by performing pairwise alignment of the hydrogenase amino acid sequences using MUSCLE v3.8.1551^75^. Finally, the distribution of pS^(site)^ and pN^(site)^ values on the hydrogenase structure were visualized using ‘anvi-display-structure’.

### Statistical analyses

Statistical analysis was carried out in R (v4.0.0). The Kruskal-Wallis rank sum test with Chi-square correction was used for comparison of pN/pS in genes among different phyla and sites. The Wilcoxon rank-sum test was used for comparison of pN/pS in genes between two groups. Linear regression was used to fit the data and predict the linear correlation between the two indexes of protein biophysical characteristics and pN/pS in genes.

## Data Availability

The newly acquired metagenomic data and MAGs has been deposited in the NCBI Sequence Read Archive (SRA) database under the accession number PRJNA630822. The protein amino acid sequences, protein structures and phylogenetic trees of hydrogenases are available at figshare (https://doi.org/10.6084/m9.figshare.30000151). All other study data are included in the article and/or supporting information.

## Acknowledgements

We appreciate the computational resources and platforms provided by the MonARCH HPC Cluster and the MASSIVE HPC facility, as well as the detailed codebase developed by Anvi’o for structure-informed microbial population genetics. We thank Mike McDonald and James Lingford for their helpful discussions and insights.

## Funding statements

This work was supported by ARC SRIEAS Grant SR200100005 Securing Antarctica’s Environmental Future and contributes to delivering the Australian Antarctic Science Decadal Strategy. C.G. was supported by an ARC Future Fellowship (FT240100502). P.M.L. and R.L. were supported by ARC DECRA Fellowships (DE250101210 and DE230100542). Y.P. was supported by the Monash Graduate Scholarship (MGS) and Monash International Tuition Scholarship (MITS) awarded by Monash University.

## Author contributions

C.G., Y.P., P.M.L., L.C.W., X.D., and S.L.C. designed this study. L.P.J., P.M.L., R.L., and C.G. conducted long-read metagenomic analysis. Y.P., L.C.W., and C.G. conducted horizontal gene transfer analyses. Y.P., X.D., and C.G. conducted selection analysis. M.J. and Y.P. conducted structural analyses. Y.P., C.G., and S.R.H. wrote the manuscript with contributions from all authors.

## Conflict of interest

The authors declare no conflict of interest.

## Notes

### Competing Interest Statement

The authors have declared no competing interest.

