## Supplementary Figures for "Widespread horizontal transfer and strong selection enhance microbial adaptation in Antarctic soils"

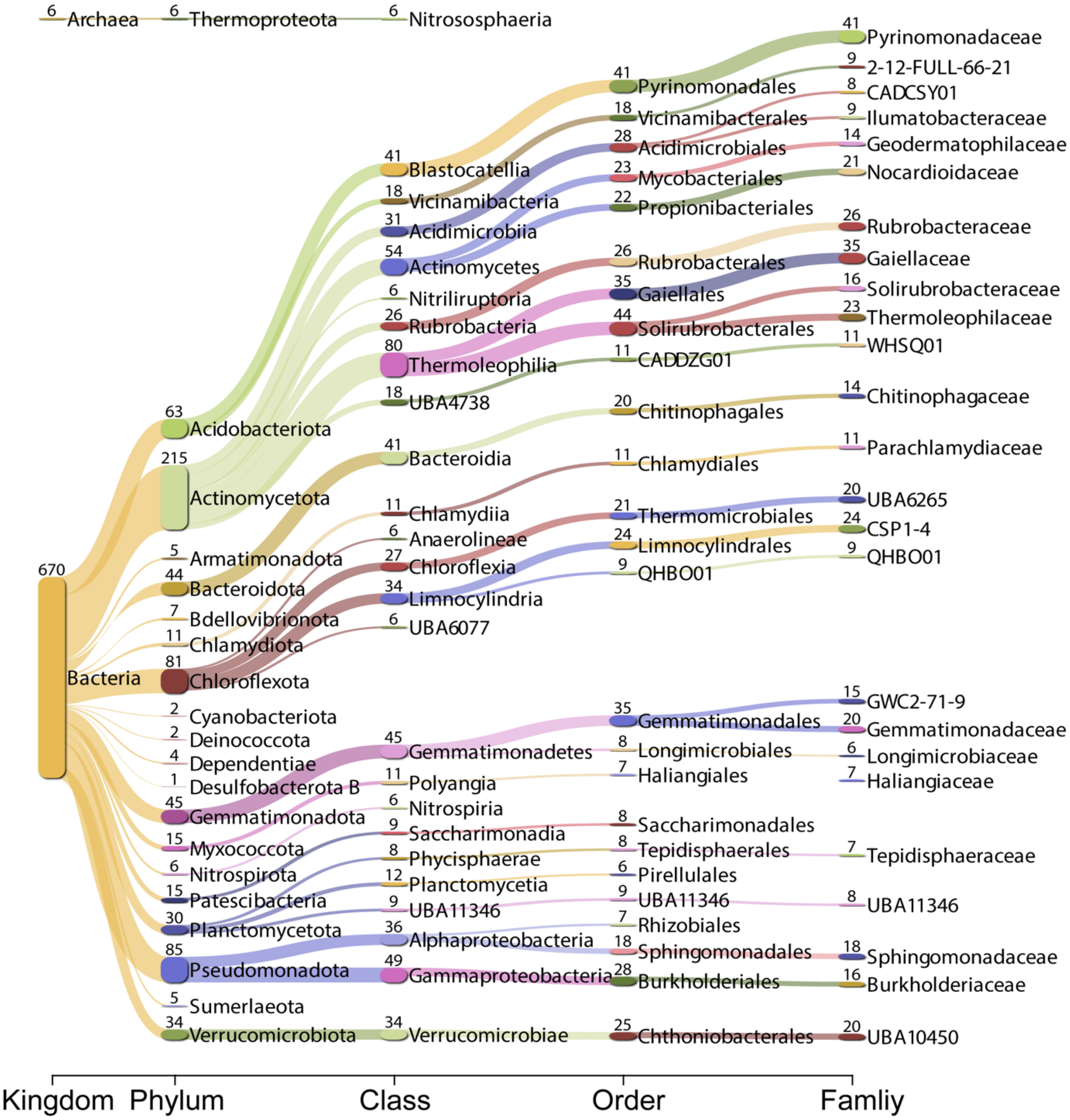


**Supplementary Figure 1. GTDB-based taxonomic classification of archaeal and bacterial MAGs recovered from the Mackay Glacier region.** The Sankey diagram illustrates the number of MAGs assigned to each lineage across different taxonomic levels. Only the top 25 taxa with the highest number of MAGs at each level are shown. Detailed statistics for 676 MAGs are provided in Supplementary Table 2.

a

| | Actinomycetota | Bacteroidota | Planctomycetota | Verrucomicrobiota | Myxococcota | Chlamydiota | Armatimonadota | Patescibacteria | Nitrospirota | Desulfobacterota_B | Dependentiae | Cyanobacteriota | Thermoproteota | Sumerlaeota | Bdellovibrionota | Deinococcota | Acidobacteriota | Gemmatimonadota | Chloroflexota | Pseudomonadota |
|---|---|---|---|---|---|---|---|---|---|---|---|---|---|---|---|---|---|---|---|---|
| Actinomycetota | 3563 | 19 | 71 | 38 | 91 | 4 | 3 | 0 | 1 | 1 | 1 | 1 | 2 | 3 | 8 | 12 | 84 | 128 | 450 | 124 |
| Bacteroidota | 28 | 333 | 9 | 11 | 3 | 2 | 0 | 1 | 1 | 0 | 0 | 0 | 0 | 0 | 1 | 0 | 27 | 14 | 8 | 19 |
| Planctomycetota | 76 | 1 | 121 | 90 | 23 | 0 | 3 | 0 | 1 | 0 | 0 | 0 | 0 | 0 | 1 | 1 | 45 | 39 | 51 | 189 |
| Verrucomicrobiota | 28 | 4 | 96 | 186 | 7 | 1 | 1 | 0 | 1 | 0 | 0 | 0 | 0 | 1 | 0 | 1 | 50 | 20 | 9 | 77 |
| Myxococcota | 166 | 6 | 27 | 9 | 45 | 0 | 1 | 0 | 1 | 0 | 0 | 0 | 0 | 1 | 0 | 0 | 67 | 35 | 25 | 52 |
| Chlamydiota | 1 | 1 | 0 | 0 | 0 | 51 | 0 | 0 | 0 | 0 | 1 | 0 | 0 | 0 | 0 | 0 | 1 | 0 | 5 | 2 |
| Armatimonadota | 9 | 4 | 3 | 6 | 1 | 0 | 2 | 0 | 0 | 0 | 0 | 0 | 0 | 1 | 0 | 0 | 30 | 5 | 7 | 21 |
| Thermoproteota | 0 | 11 | 0 | 0 | 0 | 1 | 0 | 1 | 0 | 0 | 0 | 0 | 0 | 0 | 0 | 0 | 0 | 0 | 0 | 0 |
| Patescibacteria | 0 | 1 | 0 | 0 | 0 | 0 | 0 | 5 | 0 | 0 | 0 | 0 | 0 | 0 | 0 | 0 | 4 | 2 | 1 | 1 |
| Cyanobacteriota | 1 | 0 | 0 | 0 | 0 | 0 | 0 | 0 | 0 | 0 | 0 | 0 | 0 | 0 | 0 | 0 | 5 | 0 | 0 | 1 |
| Dependentiae | 0 | 0 | 1 | 0 | 0 | 0 | 0 | 0 | 0 | 0 | 2 | 0 | 0 | 0 | 0 | 0 | 1 | 0 | 0 | 0 |
| Desulfobacterota_B | 3 | 0 | 1 | 1 | 0 | 0 | 0 | 0 | 1 | 0 | 0 | 0 | 0 | 0 | 0 | 0 | 10 | 2 | 1 | 15 |
| Nitrospirota | 4 | 2 | 2 | 2 | 0 | 0 | 0 | 0 | 0 | 0 | 0 | 0 | 0 | 0 | 0 | 0 | 13 | 4 | 3 | 5 |
| Bdellovibrionota | 12 | 1 | 0 | 1 | 1 | 0 | 0 | 0 | 1 | 0 | 0 | 0 | 0 | 0 | 0 | 0 | 0 | 1 | 1 | 1 |
| Deinococcota | 13 | 0 | 1 | 1 | 2 | 0 | 1 | 0 | 0 | 0 | 0 | 0 | 0 | 0 | 0 | 0 | 1 | 6 | 9 | 8 |
| Sumerlaeota | 7 | 0 | 7 | 5 | 1 | 0 | 0 | 0 | 0 | 0 | 0 | 0 | 0 | 0 | 0 | 0 | 1 | 7 | 6 | 9 |
| Acidobacteriota | 101 | 20 | 65 | 76 | 37 | 4 | 19 | 1 | 0 | 1 | 0 | 1 | 0 | 1 | 0 | 1 | 288 | 91 | 60 | 91 |
| Gemmatimonadota | 189 | 11 | 59 | 28 | 23 | 0 | 4 | 1 | 1 | 0 | 0 | 0 | 0 | 2 | 0 | 3 | 81 | 235 | 104 | 98 |
| Chloroflexota | 475 | 6 | 43 | 16 | 25 | 1 | 4 | 0 | 0 | 0 | 0 | 0 | 1 | 7 | 0 | 7 | 54 | 59 | 353 | 81 |
| Pseudomonadota | 142 | 9 | 94 | 73 | 23 | 2 | 7 | 1 | 4 | 2 | 0 | 0 | 0 | 7 | 4 | 2 | 102 | 67 | 72 | 522 |

Number of HGT: 0, 500, 1000, 1500, 2000, 2500, 3000, 3500

b

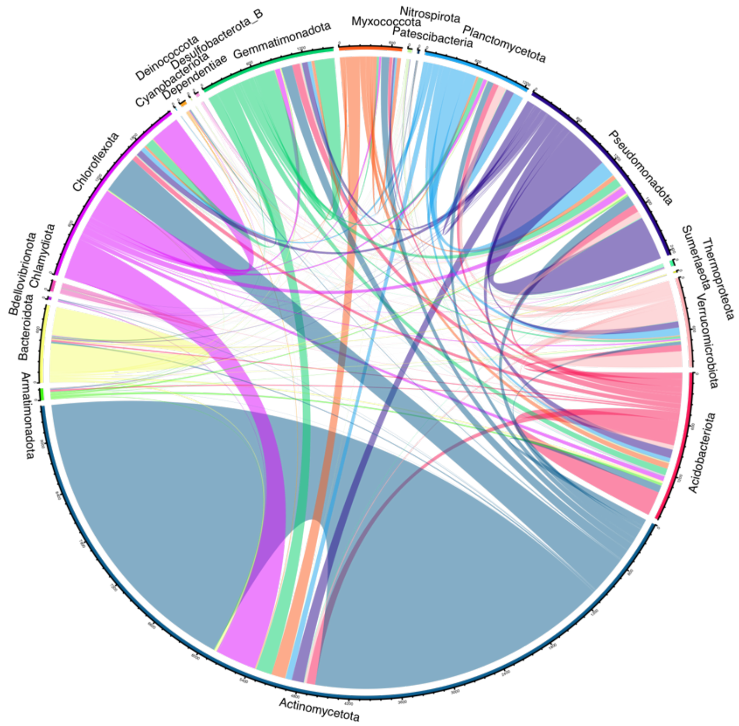


**Supplementary Figure 2. HGT events among different microbial phyla in Antarctic soils.** The (a) heatmap and (b) circular diagram display the number of inferred HGT events between pairs of microbial phyla. Detailed statistics for HGT events across 676 MAGs are provided in Supplementary Tables 3 and 4.

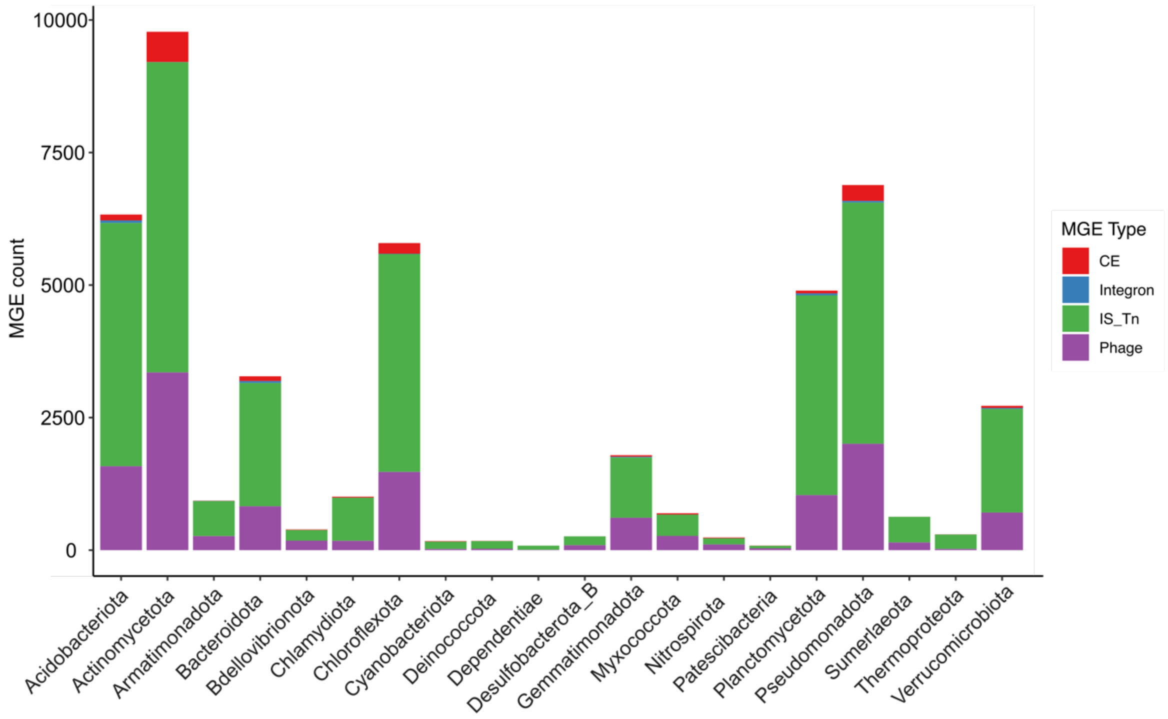


**Supplementary Figure 3. The number of MGEs across microbial phyla in Antarctic soils.** The bar plot shows the total number of MGEs identified in each phylum, with different MGE types represented by distinct colors. CE refers to conjugative elements and IS_Tn refers to insertion sequences (IS) and transposons. Detailed annotations of MGE-related genes are provided in Supplementary Table 9.

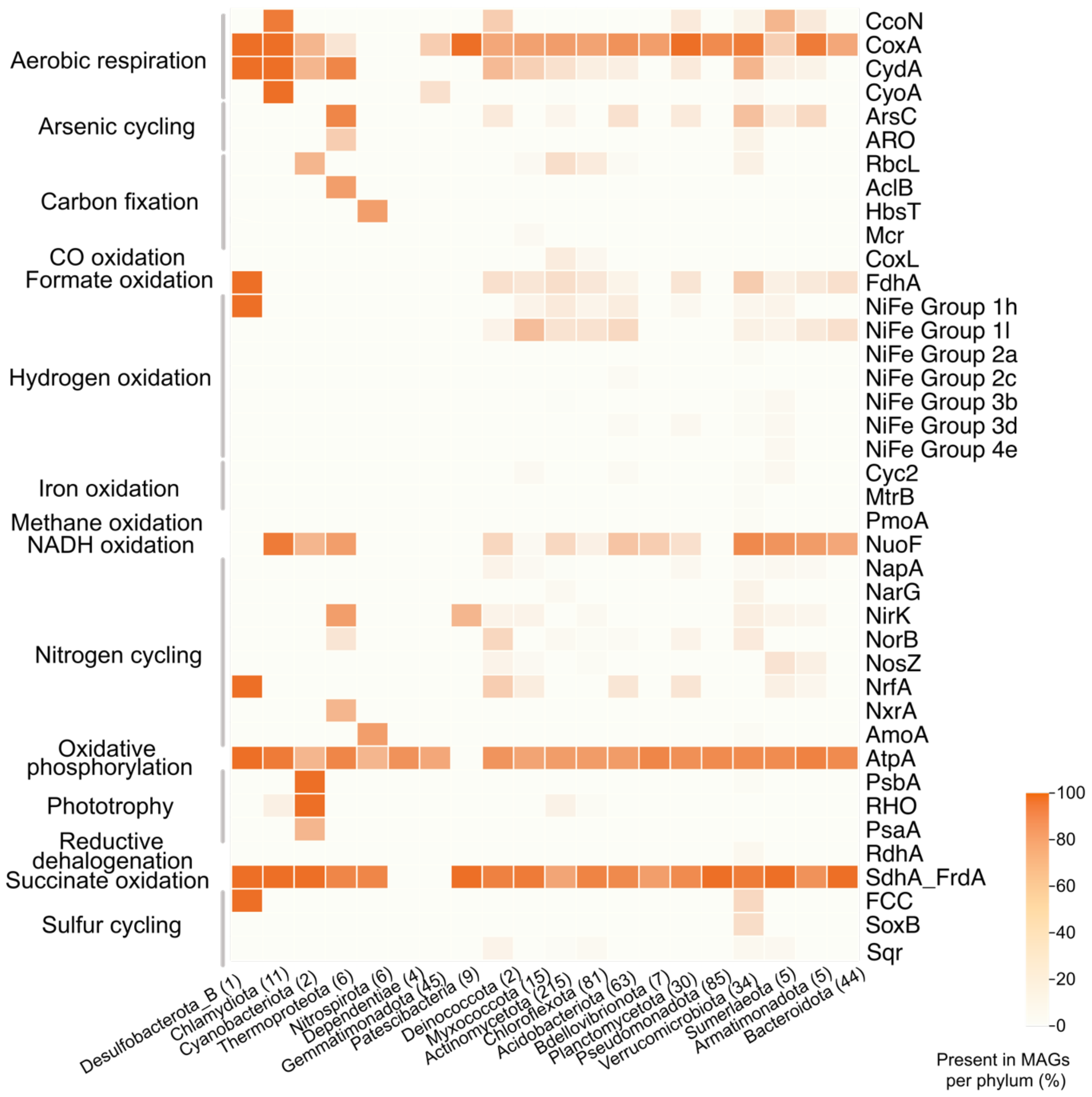


**Supplementary Figure 4. The proportion of key metabolic proteins detected across Antarctic microbial phyla.** The heatmap shows the proportion of MAGs within each microbial phylum that contain specific key metabolic proteins. Color intensity reflects the frequency of protein presence. Detailed annotations are available in Supplementary Table 8.

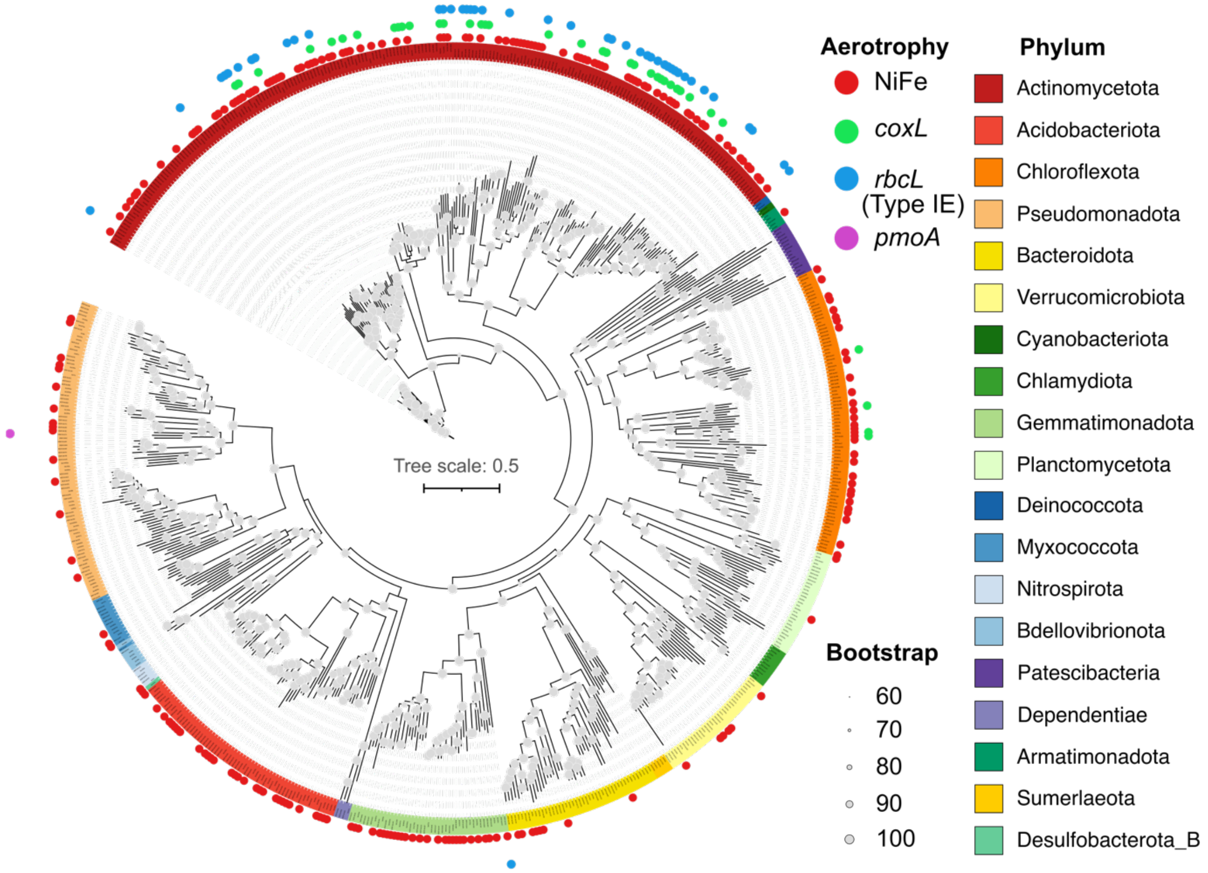


**Supplementary Figure 5. Maximum-likelihood phylogenetic tree of bacterial MAGs in Antarctic soils, annotated with phylum-level taxonomy and aerotrophy gene content.** The phylogenomic tree was inferred based on concatenation of conserved bacterial single-copy genes that were used as phylogenetic markers in GTDB-Tk, constructed using IQ-TREE. MAGs containing aerotrophy genes (group 1h and 1l [NiFe]-hydrogenase, *coxL*, type IE *rbcL*, and *pmoA*) are indicated by colored circles. Bootstrap support values (60-100%) are indicated by the size of the grey dots at each node. Scale bars represent the number of substitutions per site. Details of these MAGs and aerotrophy genes are available in Supplementary Table 8.

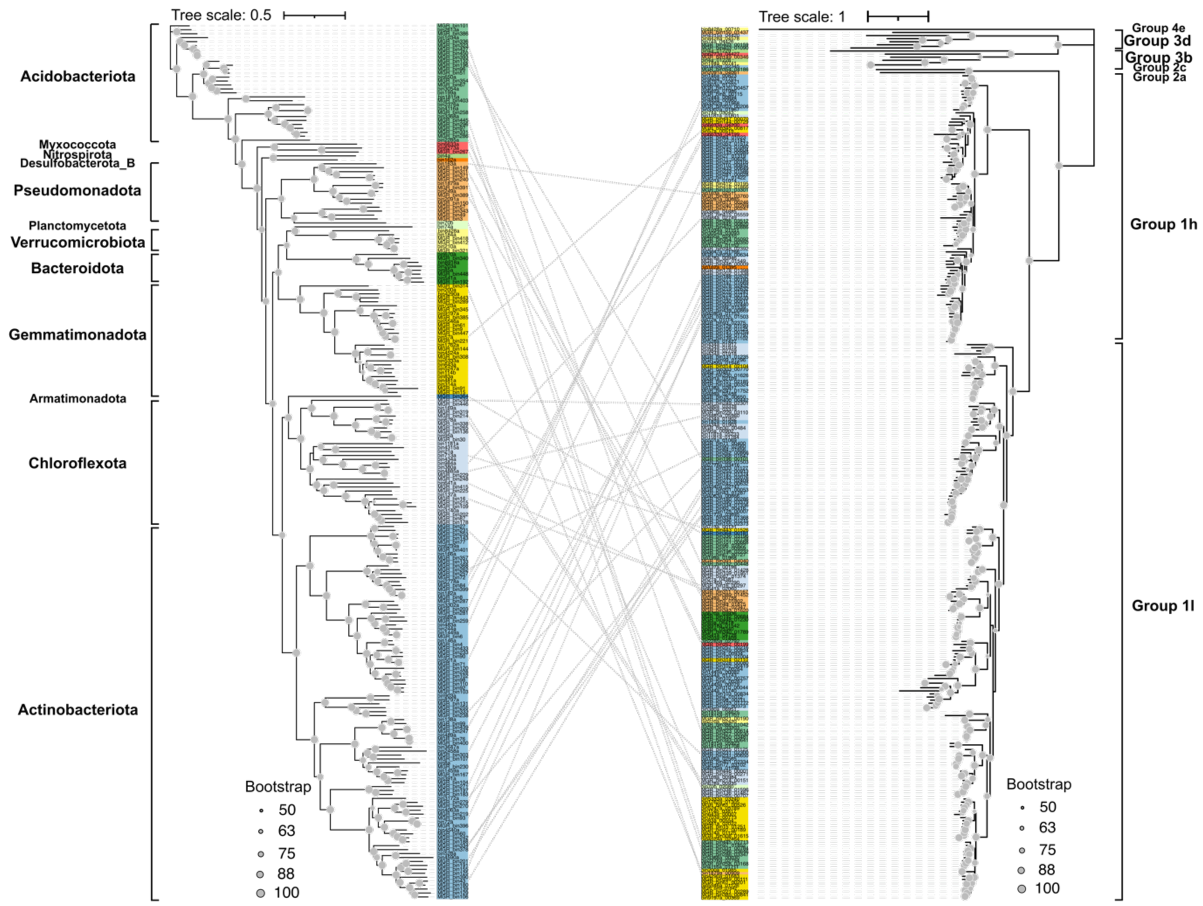


**Supplementary Figure 6. Maximum-likelihood phylogenetic trees of hydrogenase-bearing MAGs and their hydrogenase protein sequences.** Left panel shows a phylogenomic tree based on concatenated conserved marker genes from 222 bacterial MAGs encoding hydrogenases, with phylum-level taxonomy annotated. The right panel displays a phylogenetic tree of the hydrogenase protein sequences derived from the same MAGs. Hydrogenases predicted to be horizontally transferred are linked to their corresponding MAGs with dashed lines. Sequences from the same taxonomic groups are color-coded consistently across both trees. Bootstrap support values (50-100%) are indicated by the size of the grey dots at each node. Scale bars represent the number of substitutions per site. Detailed statistics for representative MAGs and hydrogenase genes are available in Tables S2 and 8.

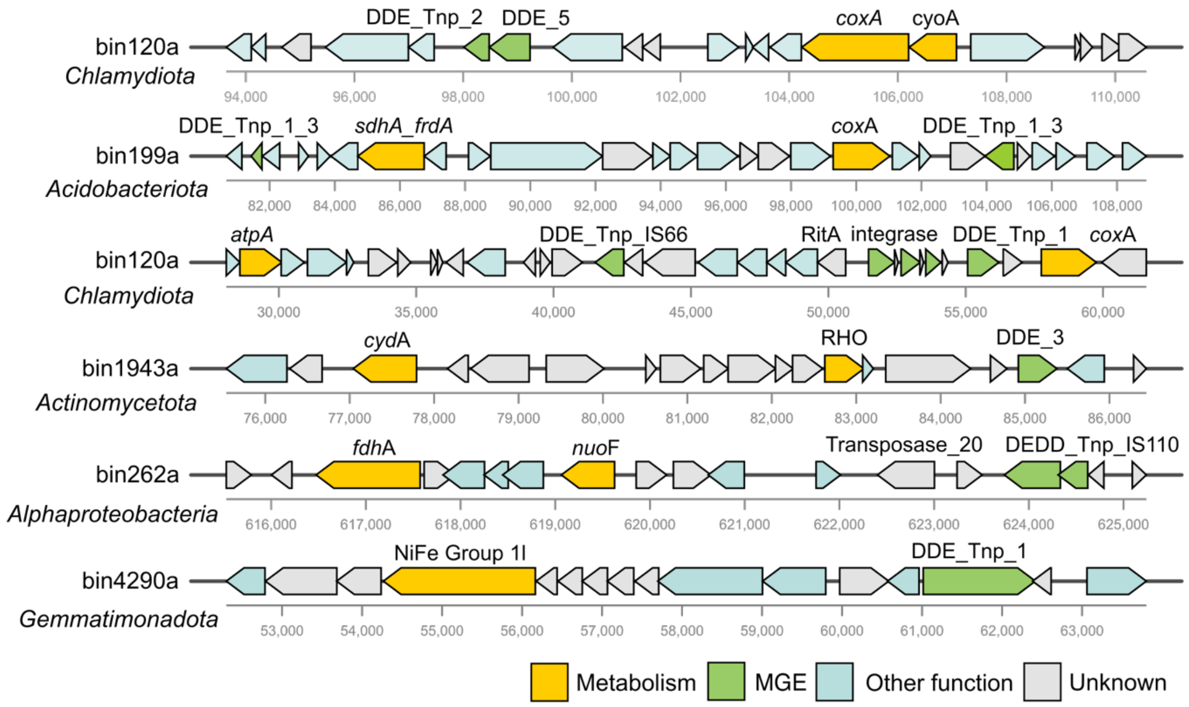


**Supplementary Figure 7. Representative gene clusters containing metabolic marker genes involved in energy conservation.** Genes are color-coded by functional category: yellow, metabolic genes; green, MGEs; blue, other functional genes; grey, unknown function. Detailed statistics for MGEs are provided in Supplementary Table 9.

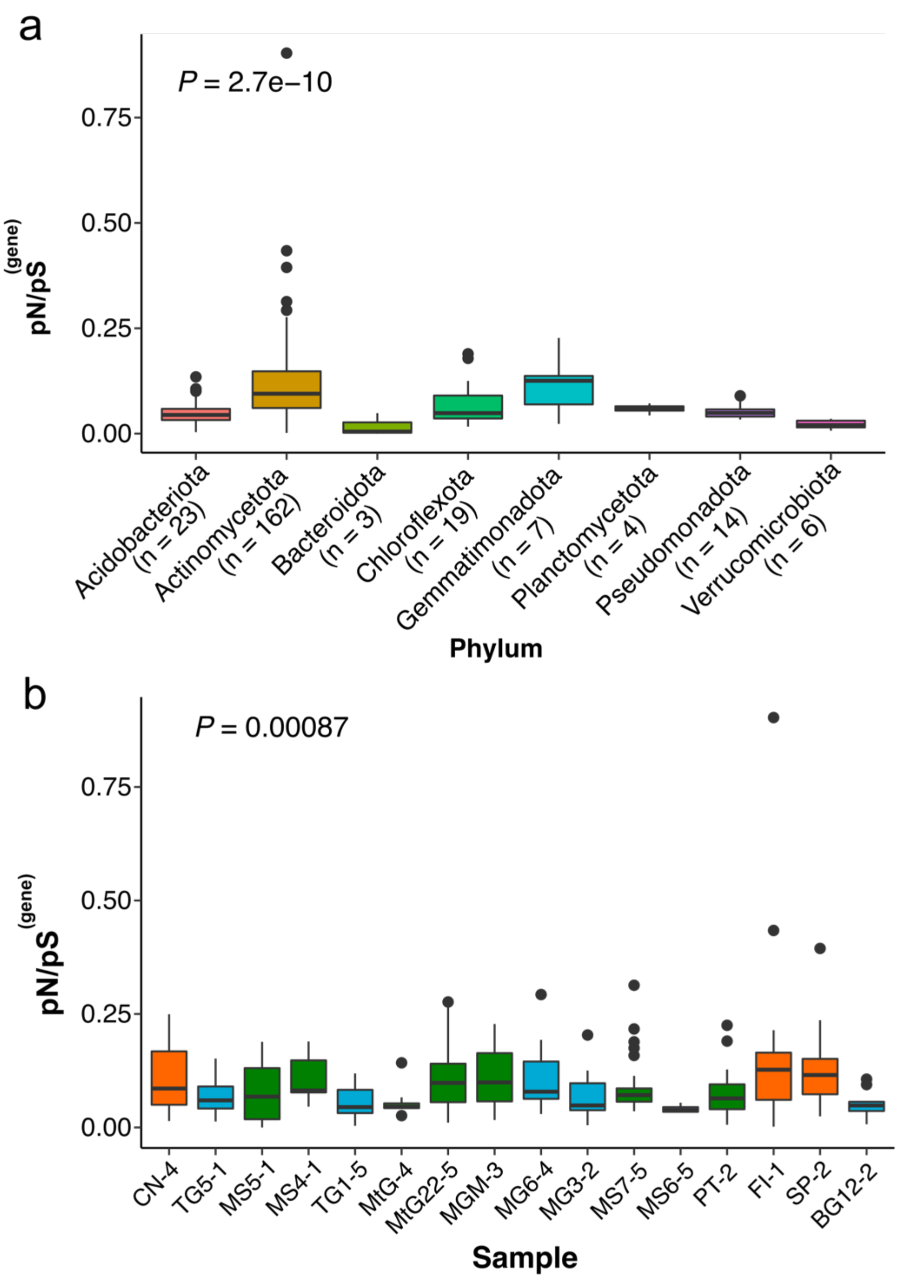


**Supplementary Figure 8. Comparison of pN/pS for hydrogenase genes in Antarctic soils.** (a) pN/pS values of hydrogenase genes across different microbial phyla (n = number of gene-sample data points analysed per phylum). (b) pN/pS values of hydrogenase genes across different sampling sites. Boxes are coloured by sample type: mountain (green), glacier (blue), and miscellaneous (orange). P-values of differences across different taxonomic groups and sample sites were calculated using Kruskal-Wallis Rank Sum test. Source data can be found in Supplementary Tables 1 and 10.

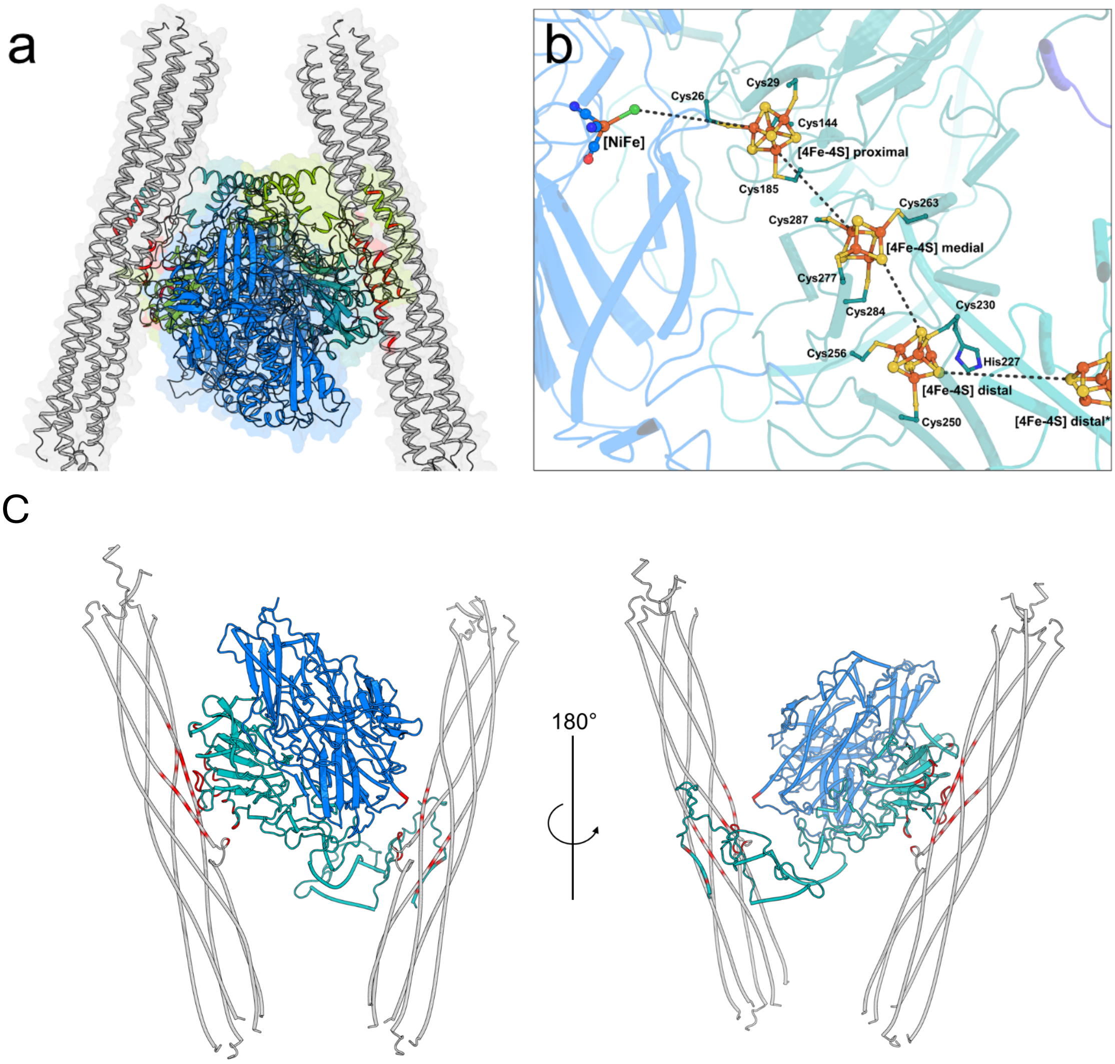


**Supplementary Figure 9.** (a) The interface of Hyl with the transmembrane helices is primarily established by HylS. Contacts involved in the oligomerization of the Hyl dimer ($(HylSU)_2$) and the two membrane tubes (HylTM1-5) are highlighted in red. (b) Electron transfer relay within one Hyl dimer. The [NiFe] active site and the [4Fe-4S] clusters are shown as balls and sticks, with the proposed electron flow from the [NiFe] active site indicated by black dashed lines. (c) Predicted contacts involved in the oligomerization of the 11 [NiFe]-hydrogenase with its membrane stalks are highlighted in red. The cartoon shows one large subunit (HylL, blue) and one small subunit (HylS, teal) and both transmembrane stalks (HylTM1-5, grey). The interface between the two stalks (HylTM1-4) and the hydrogenase is predominantly established by the small subunit.

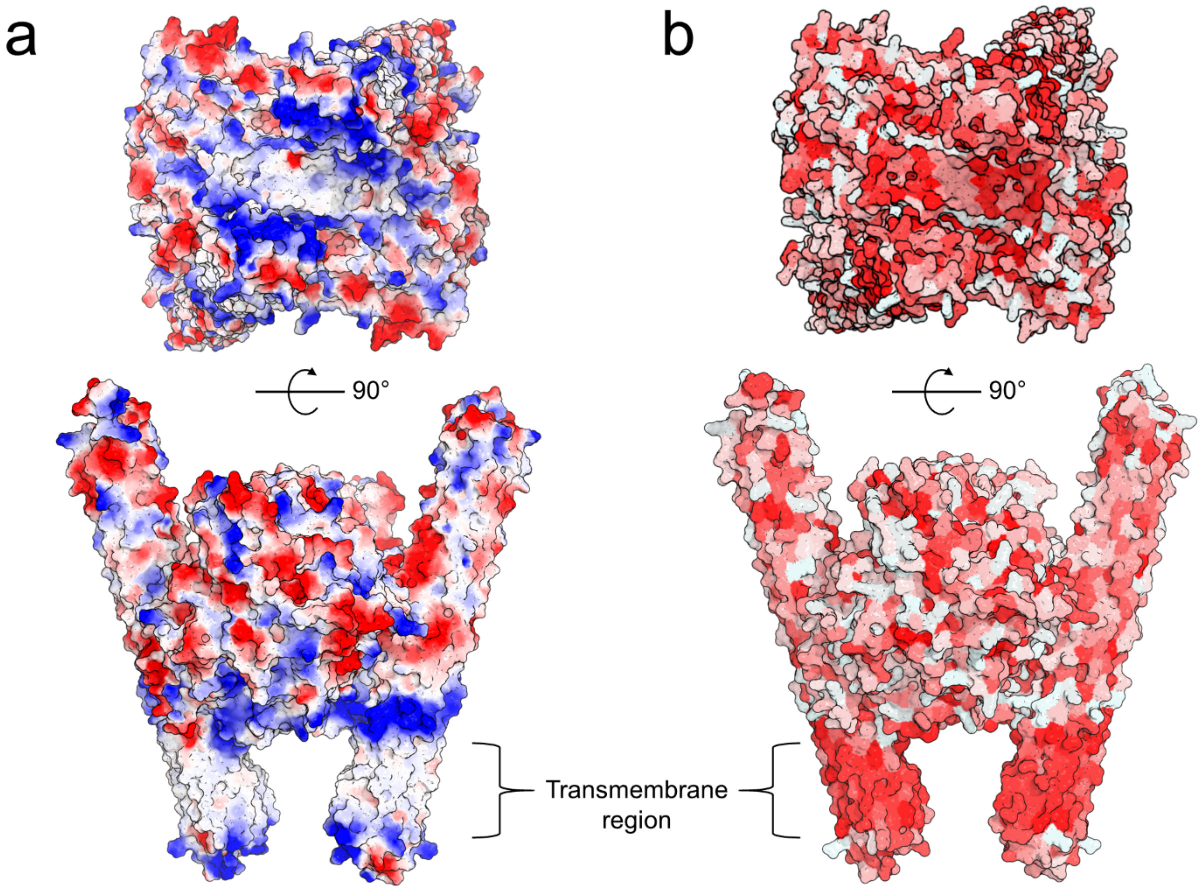


**Supplementary Figure 10. Predicted vacuum electrostatics and hydrophobicity surface analysis of the group 1l [NiFe] hydrogenase complex.** The predicted structural model of the group 1l [NiFe] hydrogenase was generated from the experimentally validated $H_2$ oxidizer *Hymenobacter roseosalivarius*. (a) Vacuum electrostatic surface analysis, shown as a gradient from red to blue for a more negative to a positive potential, respectively, with white indicating neutral surface potential. (b) Hydrophobicity surface analysis, shown in a hydrophobic gradient with non-hydrophobic amino acids in white to hydrophobic amino acids in red. The predicted structural model is available on Figshare (DOI: 10.6084/m9.figshare.30000151).

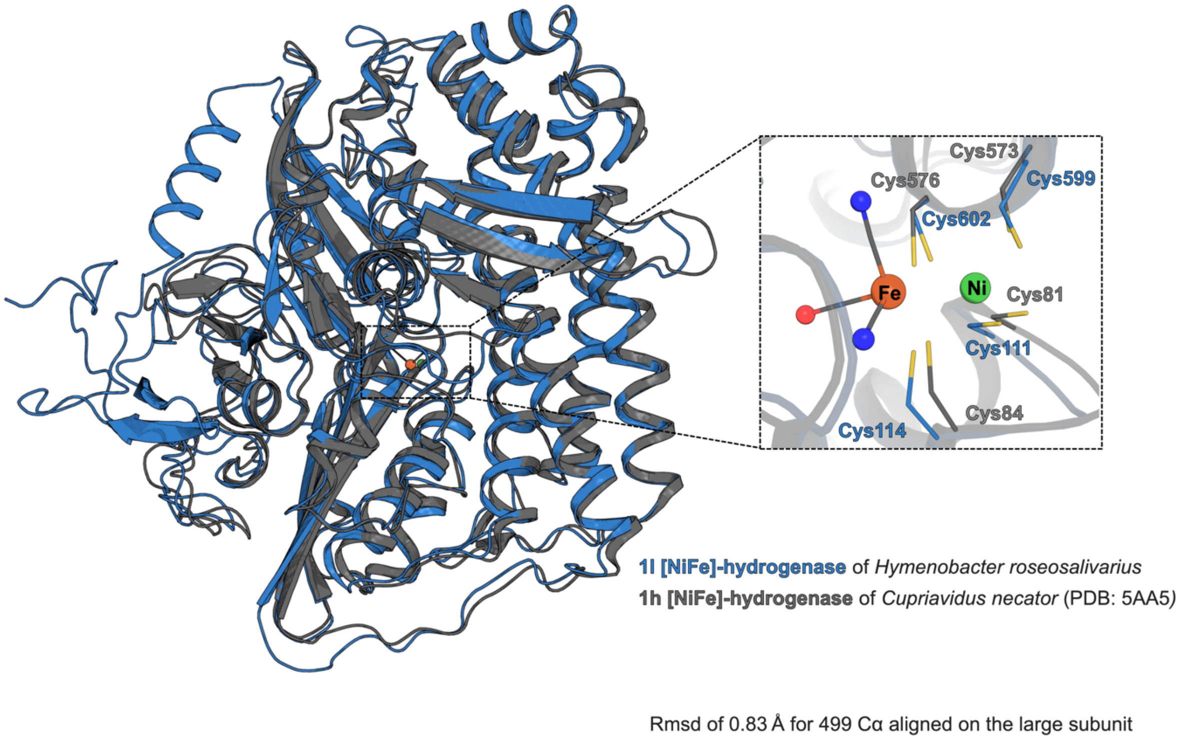


**Supplementary Figure 11. Superposition of the active sites of group 1l [NiFe] hydrogenase and the group 1h [NiFe] hydrogenase**. Structural alignment of the large subunit of group 1l [NiFe] hydrogenase from *Hymenobacter roseosalivarius* (blue) and group 1h [NiFe] hydrogenase (grey; PDB: 5AA5, rmsd of 0.83-Å for 499-Cα aligned). The [NiFe]-active site and coordinating ligands are shown in a close-up in stick and sphere representation. Carbon, nitrogen, oxygen, sulfur, iron and nickel are coloured in magenta/grey, blue, red, yellow, orange and green, respectively.

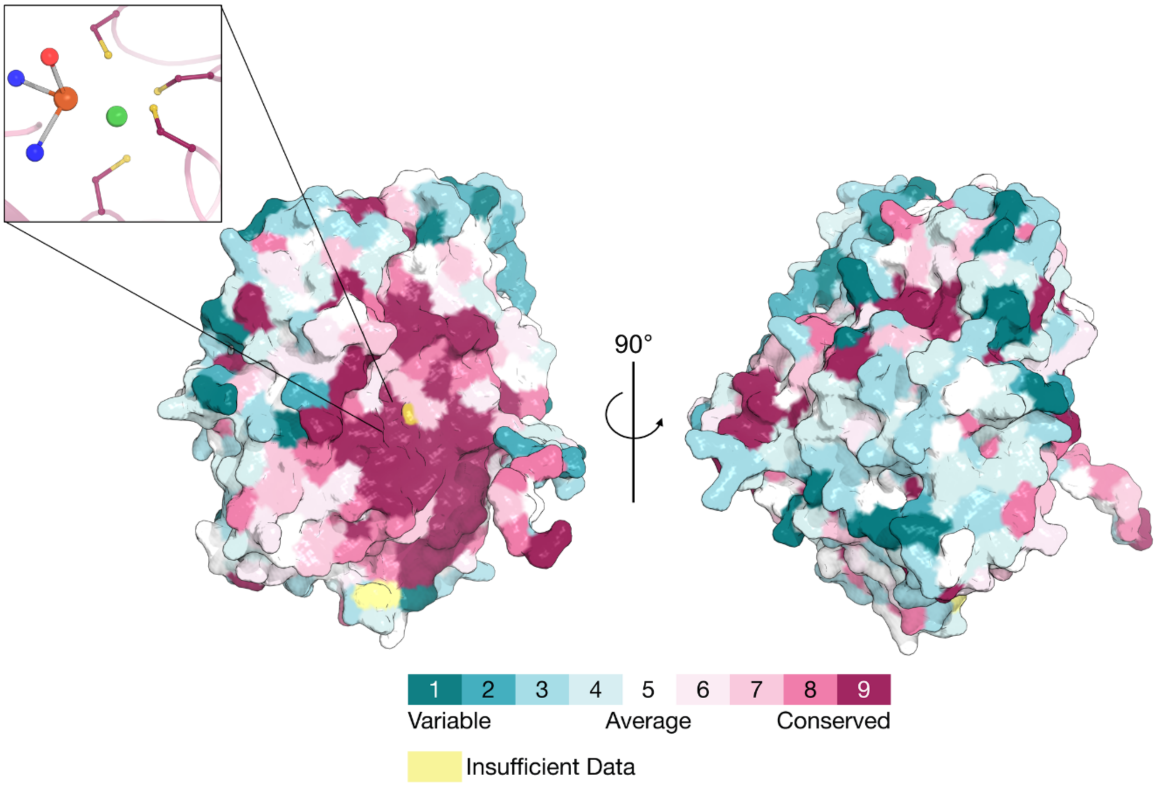


**Supplementary Figure 12. Evolutionary conservation of amino acid positions in the catalytic subunit of a group 1l [NiFe] hydrogenase protein structure based on the phylogenetic relations between homologous sequences.** The large subunit of the 1l [NiFe] structure shown in surface representation and coloured by its sequence conservation. The conservation scores are shown in a gradient from variable (turquoise), through intermediately conserved (white), to conserved positions (magenta). A close-up shows the [NiFe]-active site and the highly conserved ligating cysteine-residues, in sphere and stick representation. Position-specific conservation scores were calculated using the Bayesian method.
